# Beyond circadian: a yearlong prospective cohort study into human sleep using subcutaneously implanted electroencephalography

**DOI:** 10.1101/2025.09.14.676090

**Authors:** Tamir Avigdor, Jonas Duun-Henriksen, Esben Ahrens, Alyssa Ho, Matthew Moye, David Carlson, Birgit Frauscher, Sándor Beniczky

**Author notes:** These authors contributed equally to this work (Birgit Frauscher, Sándor Beniczky). Corresponding author: Sándor Beniczky, Address: Visby Allé 5, 4293 Dianalund, Denmark.

## Abstract

**Background:** Sleep is vital for brain health and plays a key role in many neuropsychiatric conditions. Despite its importance, sleep physiology has primarily been studied in short-term laboratory settings, limiting our ability to capture ultralong-term changes in EEG-defined sleep architecture across days, weeks, and months under real-world conditions.

**Methods:** This study leveraged a novel technology, subcutaneous electroencephalography (sqEEG) to examine ultralong-term variations in sleep. We recorded sqEEG from 20 healthy adults over a full year. We assessed intrinsic multi-day rhythms at the individual-subject level and we evaluated the effects of external factors, including season, temperature and weekends across multiple sleep metrics.

**Findings:** Early summer was characterized by shorter total sleep time and lower REM sleep duration. Weather influenced sleep: bright sunshine was associated with shorter sleep latency, whereas greater cloud cover was associated with longer sleep latency, and higher temperatures were associated with earlier sleep timing and reduced deep sleep. Weekend nights were associated with delayed sleep onset, offset, and midpoint, along with modest changes in Sunday-night sleep architecture. Social jetlag of at least one hour was present in a meaningful minority of participants, while overall sleep regularity remained high. Robust multi-day intrinsic cycles were not detected.

**Interpretation:** Continuous long-term sqEEG monitoring reveals that sleep is a dynamic process shaped by external environmental cues. Seasonal changes, temperature, and weekend schedules contribute to variations in sleep timing and duration throughout the year.

**Funding:** Filadelfia Research Fund, Duke University startup fund, and Canadian Institutes of Health Research.

## Introduction

Classic investigation of the neurophysiological underpinnings of sleep started with the invention of the electroencephalogram (EEG),(2, 3) which revealed the macrostructure of sleep.(4-6) However, the difficulty of obtaining long-term EEG recordings under natural conditions has constrained our knowledge of how internal and external factors shape sleep architecture and its long-term cyclicity. Most studies on sleep macrostructure have been restricted to sleep laboratories, where recordings are typically short-term of one or two nights and performed under controlled conditions in unfamiliar environments(7-9). These artificial settings can alter sleep patterns, limiting the generalizability of findings to typical sleep in natural settings. The first month-long study of internal sleep rhythmic outside the sleep lab dates back to 1938, and established the ∼24h sleep cycle.(10) However, such long-term studies were difficult to conduct and thus remained rare.

The recent emergence of wearable devices that can approximate sleep patterns from modalities other than EEG has enabled sleep monitoring over several months.(11, 12) While this marks a significant step beyond traditional laboratory-based approaches, these devices lack the precision for in-depth assessment of the sleep architecture, and studies spanning a full calendar year are still uncommon. When long-term studies were conducted, they found seasonal cycles of some sleep features(13-15), influenced by geographic location,(16, 17) lifestyle factors,(18, 19) and even environmental conditions within special indoors settings such as enclosed space for mars mission training. (20-23) Although non-EEG wearables have advanced the ability to monitor sleep in naturalistic settings, they lack the precision of EEG recordings, which remain the gold standard for assessing sleep macrostructure. These devices often show limited reliability(24, 25) with low Cohen’s kappa agreement rates and high variability compared to polysomnography in features such as rapid eye movement (REM) sleep (0.13-0.37) and wakefulness during the night (0.41-0.72), depending on the device used(26). This is typically attributed to their low sensitivity(24). While polysomnography (PSG) remains the gold standard for sleep staging and provides a comprehensive assessment of sleep physiology, it is generally limited to short recording periods in controlled laboratory settings. In contrast, sqEEG enables high-quality, long-term recordings in naturalistic home environments, allowing assessment of sleep over months to years. Although sqEEG provides only two channels compared to a full PSG, previous validation studies have demonstrated high agreement with conventional sleep staging, supporting its use for long-term monitoring of sleep architecture(27).

The investigation of multi-day endogenous sleep cycles has remained largely unexplored, except in studies focused on hormonal rhythms,(28) which themselves were rarely conducted long-term in home-settings. The recent development of ultralong-term subcutaneous EEG (sqEEG) has the potential to bridge the gap between wearable sleep monitoring and the gold standard of scalp EEG.(29) Implanted beneath the skin near the ear, sqEEG enables high-quality, long-term recordings, compatible with naturalistic, everyday sleep environments.(27, 30)

In this study, we leveraged the first ultralong-term sqEEG dataset in natural setting collected from healthy subjects to examine human sleep outside the laboratory. We analyzed a full year of nocturnal sleep recordings to investigate how internal and external factors influence sleep macrostructure. Our findings reveal that several aspects of sleep architecture exhibit intrinsic rhythmicity and are modulated by seasonal variation, weather conditions, and other contextual influences.

## Methods

### Subjects, recordings and sleep scoring

We conducted year-long nightly recordings from 20 healthy subjects (11 females; median age 28 years, range:19-62) using a sqEEG recording device(30) (24/7 EEG SubQ, UNEEG Medical A/S, Lillerød, Denmark) recruited between 2020-09-07 to 2022-07-04. An electrode with three contacts (two channels) was subcutaneously implanted with electrode locations of approximately TP9/TP10, T9/T10 and FT9/FT10 (Supplementary Fig. S1), according to the international 10/10 system dependent on implanted hemisphere. An external device was adhesively attached to the skin over the housing of the implant, which was connected through an inductive link for power transmission and data storage. Recordings were sampled at 207 Hz and bandpass filtered between 0.5 and 48 Hz. The device also has an in-device accelerometer to measure the displacement and velocity of movement. Participants wore the device for one year, with recordings available for 97.6% of nights and individual recording durations ranging from 279 to 365 days. The study was designed as a decentralized, prospective cohort study involving 25 enrolled healthy participants (Supplementary Fig. S2), with 20 individuals completing the full 12-month protocol(30). Eligible participants were adults aged 18 years or older without self-reported medical conditions or medication use known to affect sleep.

Participants underwent implantation of a two-channel sqEEG device positioned subcutaneously over the temporal region and recorded nightly sleep data at home for one year. Participants were instructed to activate the device when they got into bed (before turning off the lights) and to switch it off immediately after waking up. During the daytime, the device was turned off, and the external battery was connected to the charger, thus enabling continuous longitudinal sleep monitoring under real-world conditions. Sleep was scored using a modified version of U-Sleep(27, 31), a convolutional deep neural network originally trained on scalp EEG and validated for use with sqEEG, that allows to incorporate both sqEEG channels simultaneously in the feature dimension. This version modified(27) the input convolutions to integrate both two channel sqEEG channels and added adjusted sampling parameters, dropout, channel-position encoding, and dataset-specific preprocessing. The original and the adapted model weights have not been publicly released. The U-Sleep automatic sleep algorithm was validated against human scores. The U-Sleep net performance, measured by F1 score, was 0.90 ± 0.04 for Wake, 0.53 ± 0.07 for N1, 0.85 ± 0.04 for N2, 0.76 ± 0.07 for N3, and 0.90 ± 0.02 for REM. Weighted by test-record count, the corresponding scores were 0.91 ± 0.03, 0.53 ± 0.07, 0.86 ± 0.03, 0.77 ± 0.08, and 0.90 ± 0.02. Although the sqEEG system does not include dedicated EOG electrodes, the subcutaneous electrode is implanted from an entry point above the ear and directed anteriorly toward the orbital region. As a result, the recordings capture eye-movement-related activity that contributes to sleep-stage classification, including REM sleep detection. We analysed a total of 7,102 nights in which the device was activated at bedtime (lights turned off). Sleep staging was performed, regardless of the recording duration.

### Ethics

This study was approved by the Scientific Ethics Committee of Region Zealand, Denmark (reference number: SJ-778), Informed consent was obtained from all participants, and the study adhered to the Declaration of Helsinki.

### Sleep macrostructure

We analyzed the sleep macrostructure as derived from the automated sleep scoring using the previously validated method with 30 seconds epochs.(27, 31) Sleep parameters were determined for each night. Sleep onset was defined as the first occurrence of either three consecutive non-REM (NREM) sleep of NREM1 epochs or a single NREM 2 epoch. Sleep offset was defined as the last epoch of non-wake sleep stage. Sleep latency was calculated from the start of sqEEG (when going to bed). REM sleep latency was defined as the interval from sleep onset to the first REM epoch. Sleep efficiency was defined from the total sleep time between sleep onset and sleep offset.

We extracted the duration spent in each sleep stage: NREM1, NREM 2, NREM 3 and REM. The total sleep time (TST) was calculated as the sum of the time spent in all stages excluding wake. The proportion of each stage was calculated relative to TST. Wake after sleep onset (WASO) was defined as the total duration of wake epochs occurring between sleep onset and sleep offset. Sleep midpoint was calculated as the half time between sleep onset and sleep offset. We also computed social jetlag, defined as the difference between mid-sleep on weekend days (Friday–Saturday) and work days (Sunday–Thursday)(32), together with a sleep-corrected variant that compares weekend and work days at the sleep edge not affected by oversleeping. It uses sleep onset for people who go to bed later on weekend days, and sleep offset for people whose work-day wake-up is cut short, so that catch-up sleep is not mistaken for circadian misalignment, and a sleep-regularity index(33) (see below). Derived metrics were screened for values falling more than three standard deviations from each subject’s own mean. When a value exceeded this threshold, the corresponding night was inspected to determine whether an identifiable error was present, such as a misdetected bedtime, sleep onset, or sleep offset marker. If a clear and correctable cause was identified, a manual correction was made before inclusion in the analyses.

### Meteorological data

Meteorological data were obtained from the Danish Meteorological Institute, which provides publicly accessible, high-resolution environmental records for the country as a whole. We extracted daily measurements corresponding to the study period. The curated weather variables included average daily temperature (°C), hourly temperature (°C), total duration of bright sunlight (hours), mean cloud cover (the percentage of the sky covered by clouds), and total daily precipitation (millimeters). All variables were inspected for completeness and consistency. The resulting dataset was temporally aligned with the nightly sleep recordings to enable day-by-day comparisons between environmental conditions and sleep macrostructure. All data were collected in Denmark, thereby providing a relatively similar environmental context for all participants.

### Measurements organization and alignment

Each subject’s data from the sqEEG was matched with their corresponding wearable device recordings and weather observations, based on the exact dates of the nights measured. To assess seasonal effects, data from all subjects were pooled and aligned and binned for every month of the year, irrespective of the specific calendar year. Nights with missing data for a given subject were excluded from the calculations. When examining nights of excessive heat (≥20 °C)(34), we extracted three nights before and after each heat event for all subjects who had a recorded night on that date. The 20°C threshold refers to the maximum temperature and was chosen because the study was conducted in Denmark, where most homes lack air conditioning; under these conditions, nights at or above 20°C are uncommon and represent meaningful heat stress for sleep. Warm temperatures in Denmark (16± 6°C) were associated with worse medical outcomes(34). For weekend effects, subjects’ data were aligned according to the day of the week and pooled together. For an exploratory analysis of the effects of the menstrual cycle, data were aligned by capturing the 14 days preceding and the 13 days following the onset of each menstrual cycle for every month where data were available.

To examine ultralong-term internal cyclicity, we tested a range of candidate cycle periods from 8 to 60 days. For each candidate period, the full recording year was divided into repeating windows of that length, and nights falling at the same relative position across all windows were compared to assess whether the sleep parameter recurred consistently at a particular point within the cycle. We ran this scan two ways for each subject and sleep parameter: once on the raw nightly values (centered on that subject’s own median), and once after removing external factors that could otherwise introduce their own periodic structure into the scan. For the second version, we first estimated the within-subject effects of seasonality, weekday, and excessively hot nights by ordinary least squares regression. Seasonality was modeled using day-of-year harmonics, weekday effects were modeled by day of the week, and heat effects were modeled using the hot-night indicator. The model residual, a zero-mean deviation from the subject’s typical level, was carried forward into the cycle test in place of the raw value. The circular intrinsic-rhythm analysis was performed identically on both series, letting the two be compared directly, with the detrended version’s cycles specifically reflecting rhythms not explained by seasonality, weekday structure, or hot nights.

For a candidate period P, scanned from 8 to 60 days, each night t was assigned a circular phase according to its position within the cycle,

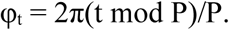

Here φₜ is the phase of night t expressed as an angle; t is the night, counted in days from the start of the recording; P is the candidate cycle period in days; (t mod P) is the position of night t within the current cycle; and the factor 2π/P maps that position onto a full circle, from 0 to 2π radians, so that nights exactly one period apart receive the same phase.

We used the phase-locking value (PLV) as a circular-timing estimate of rhythm strength, rather than a conventional t-statistic designed for linear group differences, mirroring methods used by other groups to detect multi-day cycles in EEG-derived features in epilepsy (Karoly et al., 2021). For example, in the 30-day cycle, day 1 and day 31 were both treated as the same phase (day 1) and overlaid to assess periodic patterns.

Each night’s contribution was weighted by the size of its own residual (or, in the raw version, its own centered value) at that night’s phase, so that nights with larger deviations count for more toward the resultant and nights close to the subject’s typical level count for less:

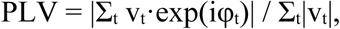

where vₜ is night t’s weight (residual or centered value) and exp(iφₜ) is the unit vector pointing in direction φₜ. PLV ranges from 0, when deviations occur at random phases and the cycle exerts no hold on the data, to 1, when the largest deviations recur at the same phase on every cycle.

Because the resultant vector is weighted by each night’s own residual rather than built from unit vectors, the classical Rayleigh test does not apply directly: its null distribution assumes unweighted vectors and is no longer valid once each vector is scaled by its own residual magnitude. We therefore used a weighted generalization of the Rayleigh test: under the null hypothesis that phase is unrelated to residual magnitude the resultant’s real and imaginary parts are asymptotically normal with variance proportional to Σₜvₜ² rather than to the number of nights; the test statistic z = 2|R|²/(τ·Σₜvₜ²), where R = Σₜvₜ·exp(iφₜ), was compared against its asymptotic chi-square null distribution, was compared against its asymptotic chi-square null distribution (p = exp(−z/2)). The factor τ ≥ 1 inflates the null variance to account for night-to-night autocorrelation in vₜ (τ = (1+ρ)/(1−ρ), from the series’ own lag-1 autocorrelation ρ; ρ can in principle range from −1 to 1, but τ is floored at 1 so that negative autocorrelation is never used to sharpen the test, and only positive autocorrelation ever inflates it). Because each subject’s series spans several hundred nights, the effective sample size n/τ remains large even when τ is not small, which is what justifies treating the resultant’s components as asymptotically normal rather than relying on an exact finite-sample distribution; this keeps a skewed or serially correlated sleep parameter, such as sleep latency, from being read as phase-locking when it is not.

The phase-locking value ranges from 0, when the highs and lows fall at random phases and the cycle exerts no hold on the data, to 1, when they recur at the same phase on every cycle; higher values therefore indicate stronger adherence to the rhythm. The Rayleigh significance and the phase-locking adherence are reported together, and the prevalence of significant cycles is summarised across subjects.(35-37). To apply a more stringent criterion and focus only on meaningful intrinsic cycles, we considered a cycle to be relevant only when the PLV exceeded 0.1. This threshold is consistent with values commonly used in biological research to define meaningful rhythmicity, where PLV values in the range of 0.1 to 0.3 are typically considered indicative of a relevant cycle, as has been shown for example in studies of multi-day seizure cycles in epilepsy(35, 36). For female participants, menstruation onset was self-reported using an Apple® Watch device 3, and the reported start date was used to align sleep parameters with the menstrual cycle.

**Figure 1.**
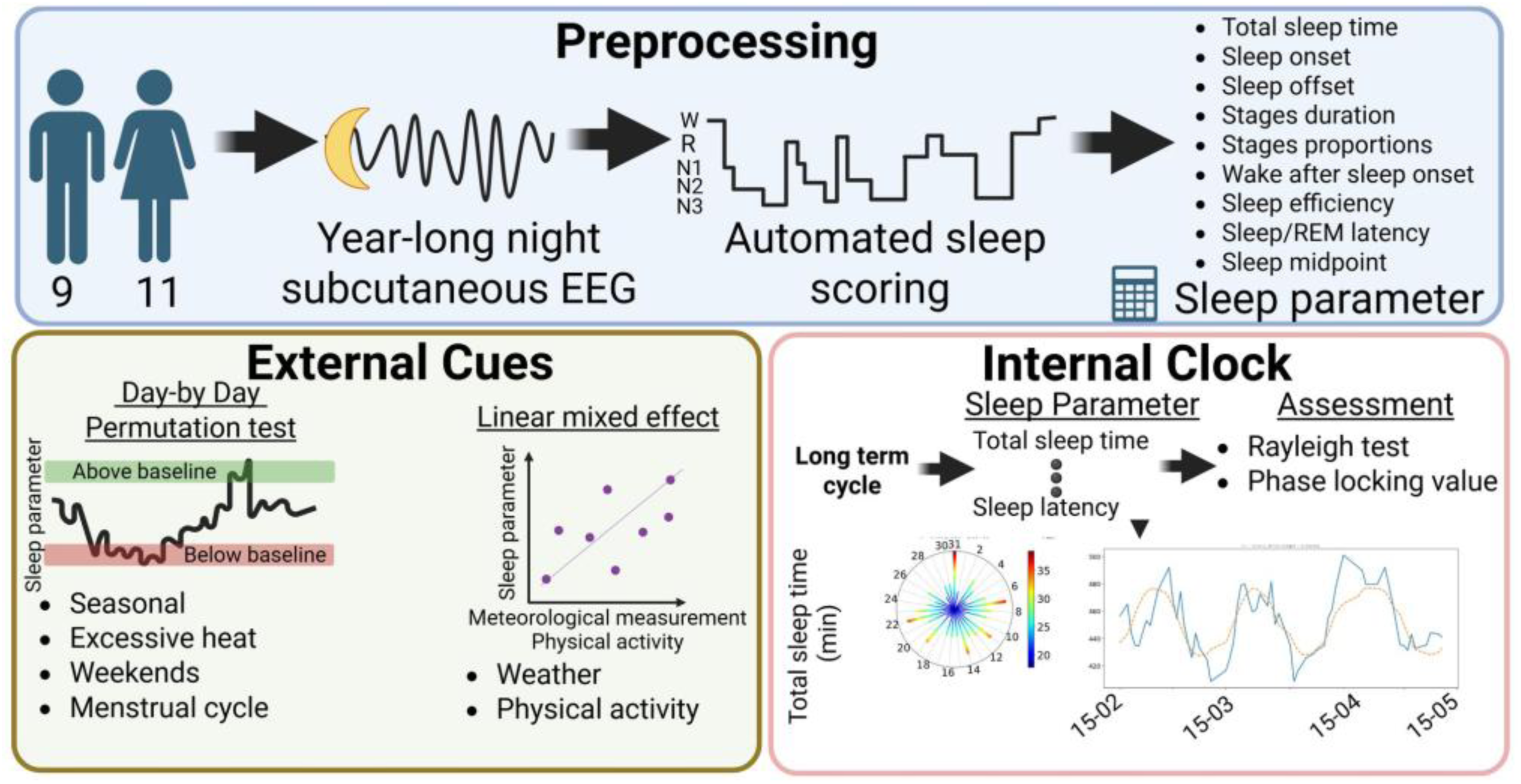
Methodological workflow. Each participant’s nightly EEG was pre-processed, automatically sleep-scored, and converted into a matrix of sleep-parameter features; the analysis was then split into two complementary tracks. Intrinsic rhythmicity was assessed by testing every sleep parameter for subject-specific cycles and, when significant, quantifying their strength with the phase-locking value. External cues modulation was assessed using permutation tests for each sleep parameter, considering season, day of the week, time of year, menstrual phase. Daily weather conditions recorded during the preceding day were modelled using linear mixed-effects models with sleep parameters accounting for the month of the year.

### Statistics

#### Longitudinal analysis

The seasonal, hot-night, day-of-week, menstrual and cohort-cycle analyses shared a single inferential procedure: a non-parametric, cluster-based permutation test (38).). It was applied separately to each sleep parameter and to each ordered axis, namely the calendar months, the nights surrounding a hot night, the days of the week, and the phases of the menstrual cycle. The test is non-parametric in the sense that it assumes no particular distribution for the data and instead derives the reference distribution of its statistic empirically, by repeatedly permuting the observed values. It was carried out in the stages described below. Each participant’s nightly series was expressed as deviations from the analysis-specific baseline, removing stable between-subject differences. At each position along the axis, we calculated a robust, median-based one-sample statistic: the across-subject median deviation relative to its dispersion. Its sign indicated the direction of change, while its magnitude reflected the strength and consistency of the deviation.

Adjacent positions with the same sign that exceeded the cluster-forming threshold, defined as the 99.5th percentile of the permutation null distribution, were grouped into clusters. Cluster mass combined the extent and strength of the deviation, favoring consistent effects spanning neighboring positions. Statistical significance was assessed using 10,000 sign-flip permutations. In each permutation, every participant’s entire deviation profile was multiplied by either +1 or −1. This preserved within-subject correlation across the axis while randomizing the overall direction of change. The group statistics and clusters were then recomputed, and the largest cluster mass was retained. An observed cluster’s p-value was the proportion of permutations with a maximum cluster mass equal to or larger than its own. Clusters with p < 0.05 were considered significant. Comparing each cluster with the largest cluster occurring anywhere under the null controlled the family-wise error rate across the axis.

#### Choice of baseline

##### Seasonal trend - whole period baseline

Each person’s baseline was their typical level across the entire recording period (the median over the whole year). Using the full year makes the reference as stable as the data allow, because it draws on every available night. Each month is then read as a departure from the person’s annual norm, which is precisely the seasonal question. A shorter, season-local baseline would absorb part of the seasonal pattern under test and shrink the very effect of interest.

##### Hot nights - two weeks prior to the hot night

Hot nights are dates in which the temperature exceeds 20 °C Because the sleep recording is keyed to the following morning, each hot date is advanced by one day to the night-key of the morning after the sleep night it actually affects, which is taken as day 0. The displayed and tested window spans several nights before and after day 0, so that the nights leading up to the hot night are themselves examined. The baseline is the 14-day window lying immediately before the displayed pre-event nights that is, it ends a few nights before the hot night and does not include the nights shortly preceding it. Each night in the window is expressed as a within-subject median deviation from this 14-day baseline (after the parameter has been residualised within subject and day of week). Fourteen days is long enough to estimate the subject’s current typical level stably while remaining local in time, so the hot-night response is measured against a season-appropriate reference rather than against the whole-year average,

##### Weekend - Monday to Thursday baseline

The axis is the seven days of the week. Each person’s baseline is their Monday through Thursday nights, the most routine work nights, which are free of weekend effects. Each remaining night is then expressed as a within-person median deviation from this baseline. The nights of primary interest are Friday and Saturday, when sleep onset and offset typically shift later, and Sunday, which represents the transition night during which sleep timing begins shifting back toward the work-week schedule. Using Monday through Thursday as the fixed baseline isolates the timing and efficiency changes seen on weekend nights and the Sunday transition, and the cluster permutation test then identifies which of these nights differ significantly from that norm

##### Menstrual cycle - whole period baseline

Subjects with at least 6 recorded onsets were included. Each parameter was residualised within subject on the annual cycle and aligned by cycle phase over a 28-day window (14 days before to 14 days after onset). As in the seasonal analysis, the baseline was the subject’s median across the whole cycle window, so each phase is a deviation from the cycle norm; the phases were tested with the cluster permutation of over the 28-phase axis treated as circular, so a perimenstrual run wrapping across onset forms a single cluster. Each person’s baseline was their average level across the whole 28-day cycle window (after removing the time of year), so that each phase is read as a departure from the cycle norm, in the same spirit as the seasonal analysis. Note that this analysis was exploratory and thus not FDR corrected, and is based on a limited cohort.

#### Weather associations

The relationship between each daily weather variable and each sleep parameter was estimated with a linear mixed-effects model fitted across all subjects and nights jointly. For a given weather–sleep pair the model was

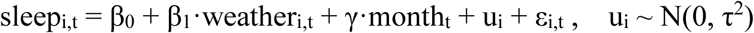

where i indexes subject and t the night. The fixed effect of interest is β₁, the slope of the sleep parameter on the weather variable; calendar month enters as a fixed covariate γ (a categorical term with twelve levels); and a random intercept is fitted for each subject.

Because sleep and weather each track the seasons independently, a raw correlation between them would partly reflect that shared annual rhythm rather than a genuine night-to-night relationship. Accounting for month of the year strips out this overlap, isolating the direct association: β₁ is then estimated from what remains once each month’s mean has been absorbed by γ, the within-month, deseasonalised association. The subject random intercept additionally gives every subject their own baseline level, so each subject acts as their own control and stable between-subject differences cannot masquerade as a weather effect; treating the nights as nested within subjects also accounts for the correlation.

#### Intrinsic multi-day cycles

We looked for rhythms with periods of 8 to 60 days, separately for each subject and sleep parameter, using circular statistics. Before testing, we removed co-variants patterns that are not the rhythm of interest: we estimated each subject’s seasonal effect (day-of-year terms), weekday effect (day of the week), and hot-night effect (the hot-night marker) and subtracted them, leaving an adjusted residual series. For each candidate period we assigned every night a phase within that cycle and used a Rayleigh test(39) to determine whether the series groups at a preferred phase rather than spreading evenly. We used the PLV to quantify how strongly the sleep parameter adheres to that rhythm(35, 36) across the whole recording: it measures how consistently peaks and troughs recur at the same phase of the cycle across the recording period. It runs from 0, when the deviations fall at random phases and the rhythm has no real hold on the data, to 1, when they land at the same phase on every cycle, indicating a stable rhythm the parameter follows throughout the year. To apply a more stringent criterion and focus only on meaningful intrinsic cycles, we considered a cycle to be relevant only when the PLV exceeded 0.1.

#### Principal component analysis

To characterize shared variation among the 11 nightly sleep metrics, each metric was z standardized within participant, and only complete nights were retained. Pooling these values across participants and nights captured common night-to-night covariation rather than between-subject differences. The first five principal components were interpreted from their largest loadings, such as a timing axis defined by sleep onset, offset, and midpoint, and a continuity-and-architecture axis defined by efficiency, wake after sleep onset, and sleep stages. An effect-magnitude vector was then calculated for four longitudinal drivers: season, day of week, hot nights, and intrinsic multi-day cycles. Seasonal magnitude was the largest absolute deviation between each seasonal mean and the participant’s overall mean, averaged across participants. Day-of-week magnitude was the mean absolute deviation of Friday through Sunday from the Monday-through-Thursday baseline. Hot-night magnitude was the largest absolute deviation across the post-event window, averaged across participants. Intrinsic-cycle magnitude was the phase-locking value. Weather associations were excluded. Each driver was projected onto each component by summing the absolute metric effects multiplied by their absolute component loadings. Projections were normalized to sum to one across the leading components, with the largest share identifying the driver’s dominant sleep axis. These analyses were descriptive and involved no distributional assumptions or hypothesis testing.

#### Social jetlag and sleep regularity

Circadian misalignment between the imposed work-week schedule and the timing of sleep on weekend days was quantified for each subject from the nightly sleep midpoint, the clock time midway between sleep onset and offset. Weekend days were defined as Friday and Saturday nights and work days as Sunday through Thursday nights, following the cohort’s usual work-week structure. Raw social jetlag - Computed per subject as the difference between the median sleep midpoint on weekend days and the average sleep midpoint on work days (Wittmann et al., 2006). A larger value means that sleep timing drifts later on weekend days than on work days, which is the defining feature of social jetlag. Sleep-corrected social jetlag: Part of the later weekend day midpoint can reflect recovery (catch-up) sleep rather than a genuine shift of the body clock, so a sleep-corrected variant was also computed (Jankowski, 2017): when sleep is longer on weekend days than on work days, the weekend day midpoint is moved earlier by half of that extra sleep duration before the work-day midpoint is subtracted. This removes the part of the apparent misalignment that is due simply to sleeping longer at the weekend and leaves a cleaner estimate of the underlying phase difference.

Sleep-regularity index: Sleep-timing regularity was summarised as one minus the standard deviation of the nightly sleep midpoint expressed as a fraction of its maximum possible value (12 hours), so that the index approaches one for highly regular timing and falls as night-to-night timing becomes more variable.

#### Sex and age

Given the sample size of 20 participants, including 11 females and 9 males, we did not test whether sex or age modulated each longitudinal effect at every calendar month, weekday, relative night, or cycle phase. Such point-by-point analyses would require splitting an already small cohort across many repeated comparisons and would therefore be underpowered and difficult to interpret. Instead, we assessed the overall contribution of sex and age to inter-individual differences in sleep using a subject-level analysis.

For each participant, each numeric sleep feature was first summarized across the full recording period by calculating that participant’s mean value. This produced one value per subject for each sleep parameter, using the full longitudinal record while avoiding repeated point-by-point demographic testing. Sex and age were then examined together in a joint linear model for each sleep feature. Before model fitting, each sleep feature was z-scored across subjects, age was z-scored, and sex was coded as female = 1 and male = 0. For each feature, we fitted the model:

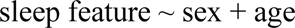

This approach allowed us to estimate the contribution of sex while adjusting for age, and the contribution of age while adjusting for sex. For each predictor, we report the standardized regression coefficient β, where positive sex β values indicate higher values in females and negative values indicate higher values in males. Positive age β values indicate that the sleep feature increases with age, whereas negative values indicate that it decreases with age.

#### Multiple-comparison correction

A Benjamini-Hochberg false-discovery-rate correction was applied for each analysis. This was applied independently within the seasonal, weekday, hot-night, weather, and intrinsic-cycle analyses. The menstrual-cycle analysis is reported at the raw (uncorrected) cluster-permutation threshold as an exploratory analysis. The following packages used for the analysis NumPy 2.1.1, pandas 2.2.2, SciPy 1.18, statsmodels 0.15.0.

## Results

These sqEEG recordings provide high-quality EEG data enabling accurate estimation of sleep across many nights of sleep. The median TST per night was 7h and 21 min, or 441 min, IQR=[391.0–484.0]. We investigated first how external cues in daily life modulated sleep as well as internal long-term sleep rhythms. We examined fluctuations in sleep macrostructure parameters, including TST, the duration and proportion of sleep stages, sleep onset and offset times, sleep latency and efficiency, sleep midpoint, and sleep disruption measured by WASO, in relation to intrinsic cyclicity and external influences

### Seasonal effects on sleep

We observed significant seasonal variation in sleep timing and architecture (Fig. 2) that deviated from the annual median. Total sleep time was lower in June, with a median reduction of 12.1 min [5.9–22.4] (p<0.05, Cliff’s d=−0.80). Sleep timing also varied across the year: sleep onset was later in July by 30.3 min [4.9–38.5] (p<0.05, d=0.50); sleep offset was earlier in September by 27.3 min [7.3–33.6] (p<0.01, d=−0.65) and later in December by 30.0 min [9.9–52.8] (p<0.05, d=0.80). Sleep architecture also showed seasonal changes: N3 duration was higher in April by 4.0 min [2.6–5.6] (p<0.05, d=0.60); REM duration was higher in February by 5.0 min [2.3–7.6] (p<0.05, d=0.70) and lower in June by 5.3 min [2.4–8.9] (p<0.001, d=−0.60) and July by 6.3 min [0.9–8.6] (p<0.001, d=−0.60). Overall, early summer was characterized by shorter sleep and lower REM duration, while sleep timing shifted later in July and December and earlier in September.

**Figure 2.**
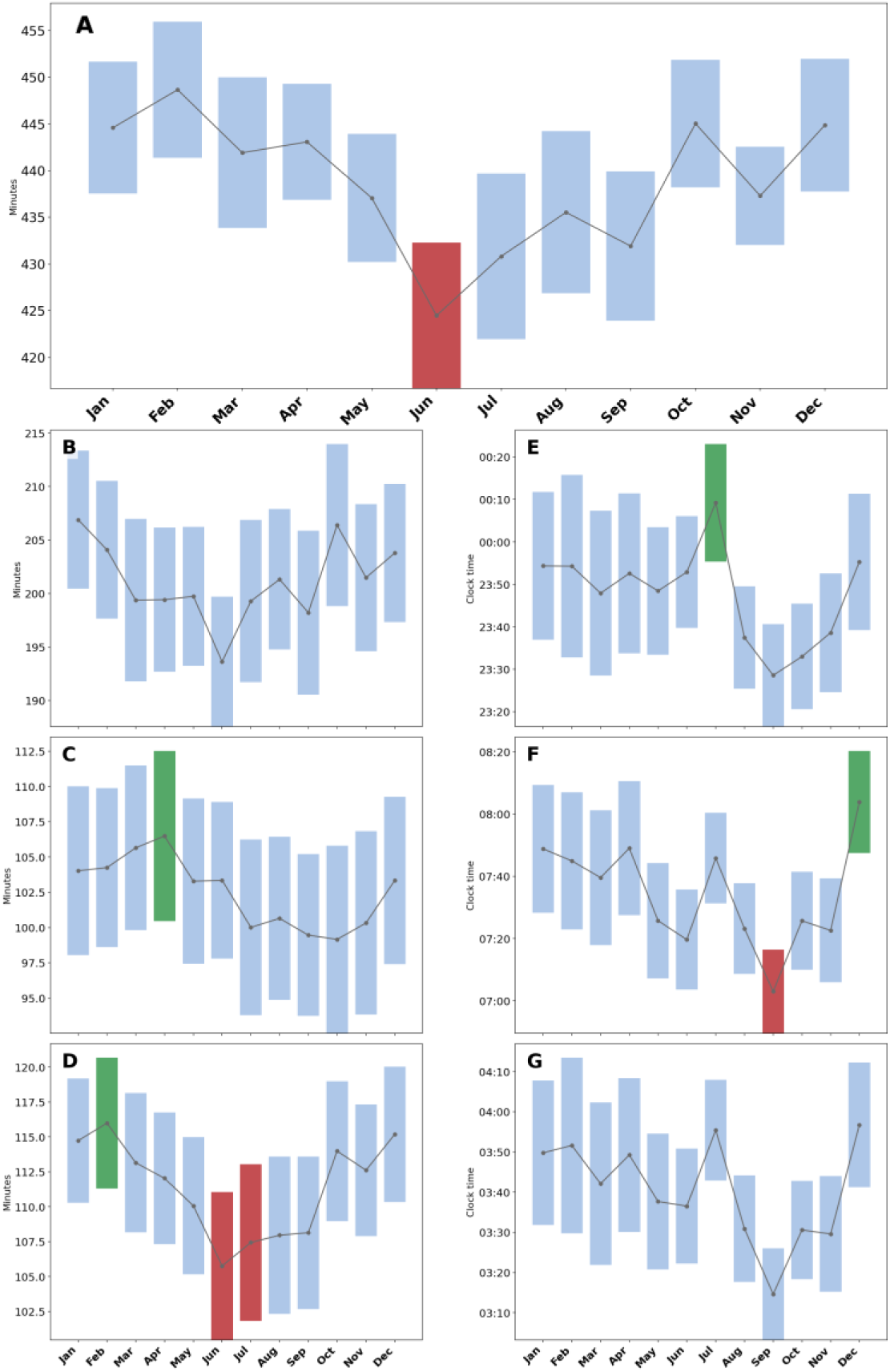
Seasonal trend in sleep. Data are pooled by calendar month across each participant’s full recorded dataset (not necessarily a single year). Each panel shows a different sleep parameter: **(A)** total sleep time (TST, minutes); **(B)** N2 sleep duration (minutes); **(C)** N3 sleep duration (minutes); **(D)** REM sleep duration (minutes); **(E)** sleep onset time; **(F)** sleep offset time; **(G)** sleep midpoint. Panels E–G are timing measures shown as clock time, anchored to the preceding local noon so that evening-to-morning values don’t wrap around midnight. For each month, the plotted point is the cohort mean and the shaded bar spans ±1 SEM, both computed from each participant’s own median value for that month. Statistical significance was assessed with a two-tailed, cluster-based sign-flip permutation test (10,000 iterations) comparing each month’s within-subject deviation against each participant’s own overall median across their full dataset, corrected for multiple comparisons via Benjamini-Hochberg false discovery rate (FDR) at a corrected α=0.05. Months with a significant increase relative to baseline after this correction are highlighted in green; months with a significant decrease are highlighted in red; non-significant months are shown in light blue.

### The effect of weather on sleep

We further explored how daily fluctuations in sleep parameters were associated with weather conditions from the prior day (Supplementary Fig. S3; standardized β±SE). Bright sunshine was associated with shorter sleep latency (β=−0.049±0.014, p<0.05), while greater cloud cover was associated with longer sleep latency (β=0.041±0.013, p<0.05). Higher daily temperatures were associated with earlier sleep timing: mean daily maximum temperature was associated with an earlier sleep offset (β=−0.090±0.025, p<0.05) and an earlier sleep midpoint (β=−0.080±0.024, p<0.05), and with reduced N3 duration, both via mean daily maximum temperature (β=−0.076±0.020, p<0.01) and mean temperature (β=−0.075±0.020, p<0.01).

On days when temperatures exceeded 20°C, the effect was more pronounced (Supplementary Fig. S4). TST decreased by a median of 18.7 min [8.1–29.4] (p<0.01, d=−0.60). WASO increased by 3.7 min [1.6–6.2] (p<0.01, d=0.80). The proportion of NREM2 increased by 0.69% [0.40–1.59] (p<0.01, d=0.60), while NREM3 duration decreased by 6.9 min [−10.3–0.3] (p<0.01, d=−0.40).

### The weekend effect

We next examined how day of week affected sleep macrostructure features (Fig. 3), using Monday–Thursday as each participant’s baseline. Sleep onset was later on Friday night by 57.6 min [40.7–81.1] (p<0.001, d=1.0) and Saturday night by 55.6 min [36.8–90.4] (p<0.001, d=1.0). Sleep offset was later on Saturday morning by 65.4 min [50.5–86.2] (p<0.001, d=0.9) and Sunday morning by 74.8 min [51.2–95.9] (p<0.001, d=1.0). Sleep midpoint was delayed on Friday night by 60.4 min [44.1–75.7] (p<0.001, d=1.0) and Saturday night by 62.9 min [40.9–81.3] (p<0.001, d=1.0). Sleep architecture showed a Sunday-night effect: the proportion of NREM2 increased by 0.95% [0.50–1.51] (p<0.001, d=0.7), while N3 duration decreased by 3.9 min [0.4–6.0] (p<0.01, d=−0.6). Overall, weekends were characterized by delayed sleep timing and modest changes in Sunday-night sleep architecture.

**Figure 3.**
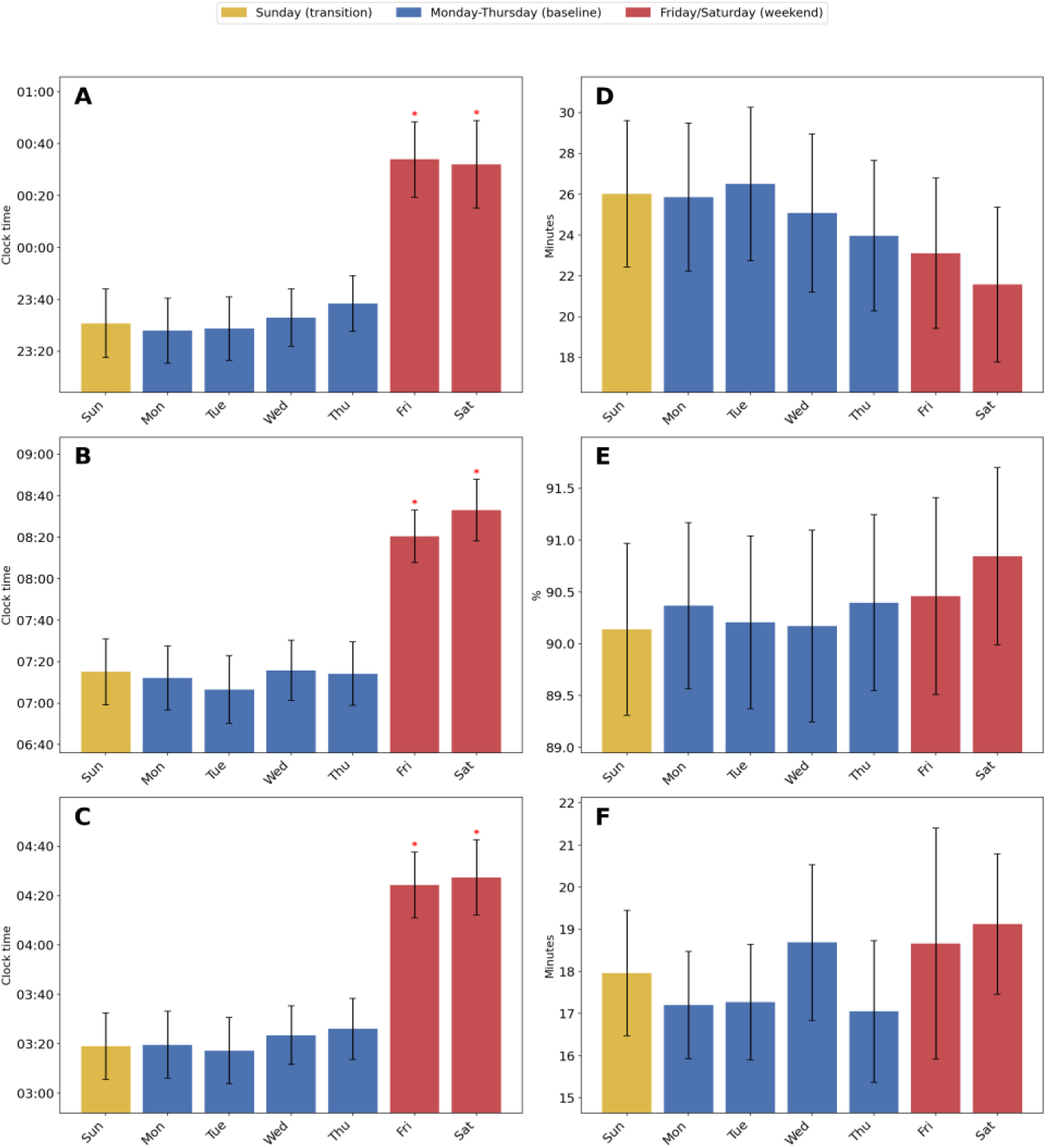
The weekend effect on sleep parameters. Panels show day-of-week variation in sleep timing and sleep continuity: **(A)** sleep onset time; **(B)** sleep offset time; **(C)** sleep midpoint; **(D)** sleep latency; **(E)** sleep efficiency; **(F)** wake after sleep onset (WASO). For each day, the plotted bar is the cohort mean (± SEM), both computed from each participant’s own median value for that day of the week. Weekday nights, Monday to Thursday, were used as the baseline and are shown in blue. Friday and Saturday nights, corresponding to the weekend period, are shown in red, and Sunday night, representing the transition back to the work week, is shown in yellow. Statistical significance was assessed with a two-tailed, cluster-based sign-flip permutation test (10,000 iterations), comparing Friday, Saturday, and Sunday individually against the weekday baseline, corrected for multiple comparisons via Benjamini-Hochberg false discovery rate (FDR) correction applied jointly across all 14 sleep metrics tracked in this analysis (not only the 6 shown here); significant effects (corrected α=0.05) are marked by asterisks. In Denmark, the weekend includes Saturday and Sunday, and therefore weekend-related sleep changes begin on Friday night and extend into it. Note that for sleep offset, the labeled night refers to the wake time on the following day.

### Intrinsic ultralong-term multi-day cycles

We did not find robust intrinsic ultralong-term multi-day sleep cycles that repeated regularly. Significance was tested using a Rayleigh test against a uniform distribution, corrected for multiple comparisons across all sleep parameters and periods tested (8–60 days), on cofactor-detrended data. For each subject-metric combination with significant cyclicity we required PLV>0.1 to be considered meaningful. Note that a PLV of 1 would indicate complete phase locking i.e., no variation in that sleep parameter on other days. After correction for multiple comparisons across all sleep parameters and accounting for external effect we did not find robust cycles, however in some instances some subject seems to partially follow a cycle (Fig. 4).

**Figure 4.**
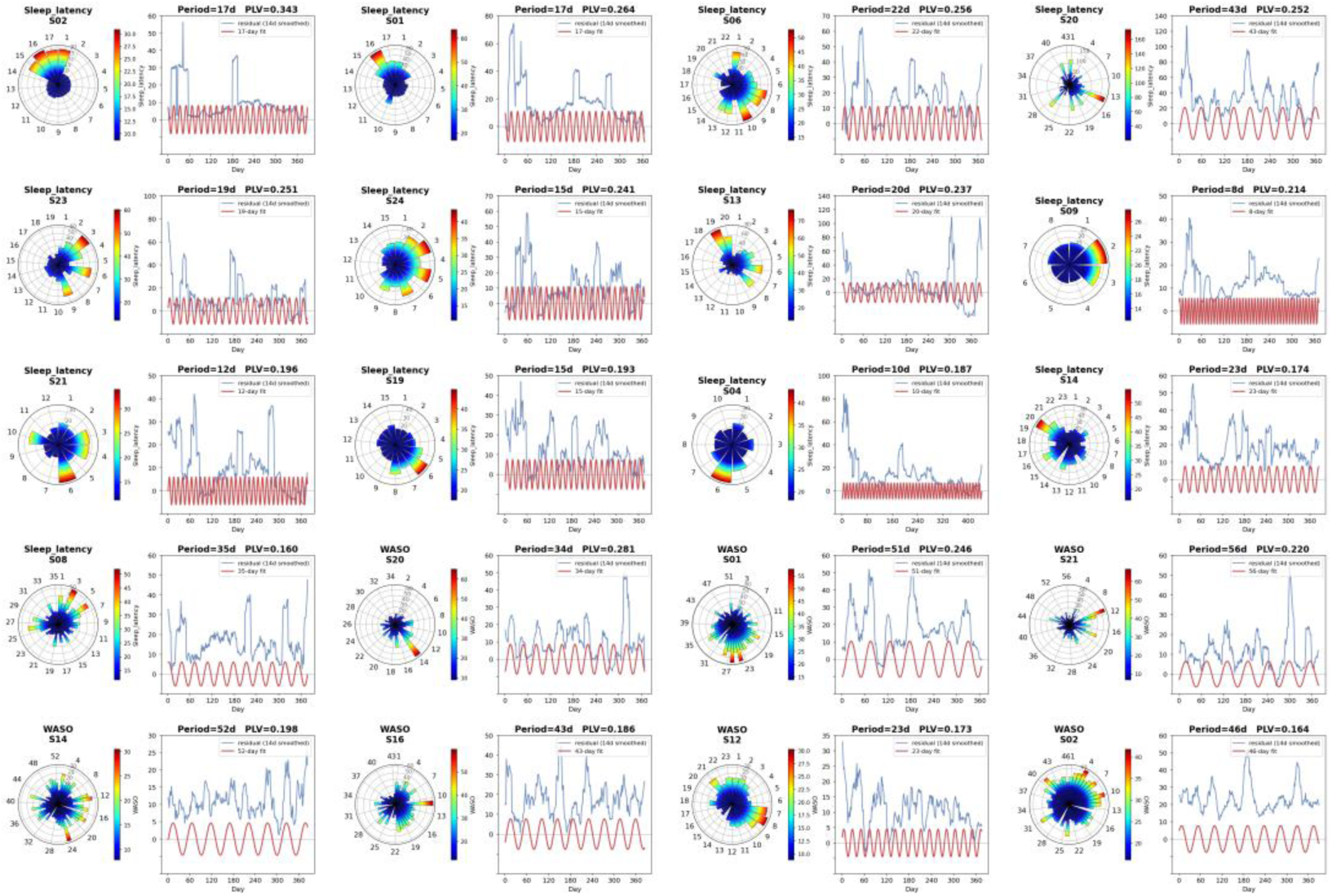
Intrinsic multi-day cycles in sleep parameters. All subject-metric combinations reaching significance are shown. A cycle was retained if its Rayleigh-test p-value survived Benjamini-Hochberg false discovery rate (FDR) correction applied across all 14 sleep metrics, all candidate periods (8–60 days), and all subjects, and if its phase-locking value (PLV) exceeded 0.1. Under this criterion, only two of the 14 tracked sleep metrics showed any significant multi-day cycles: sleep latency (13 of 20 participants) and wake after sleep onset, WASO (7 of 20 participants); total sleep time, sleep midpoint, and sleep efficiency showed none. Each panel pair shows one significant subject-metric combination. For each pair, the circular plot shows each night’s raw value of the sleep parameter, binned by its phase within that combination’s own best-fitting period and averaged within each bin; both the radial extent and the color of each wedge encode that bin’s mean value, normalized to that panel’s own value range. The adjacent line plot shows, in blue, that same series with a smoothing (14-day centered moving average). For visualization purpose only a red sinusoid is shown (least-squares fit to the period)

#### Intrinsic factors

PCA showed that the first five PCs explained 86.7% of total variance (Supplementary Fig. S5). PC1 explained 27.3% and was mainly a continuity-and-architecture axis (TST, R duration, N2 duration, sleep offset, N3 duration and sleep efficiency all loading together), while PC2 explained 23.5% and was mainly a timing axis (sleep onset, midpoint and offset). Projecting each of 4 longitudinal drivers (seasonal, day-of-week, hot night, intrinsic cycle) onto these components, a purely descriptive analysis with no significance test, showed the hot-night response had the largest projected share on every one of the first 5 components (51–65%), followed by the seasonal trend (25–31%) and the day-of-week effect (10–18%). The intrinsic-cycle driver’s projected share was negligible by comparison (<1% on every component); because that driver’s magnitude is a PLV (a 0–1 quantity) while the other three drivers’ magnitudes are raw deviations in each metric’s native units (mostly minutes), this near-zero share reflects a scale mismatch between PLV and minute-scale deviations rather than a claim that the intrinsic cycle has no structural alignment with these components; it simply isn’t directly comparable to the other three drivers’ shares as computed here.

We also performed an exploratory analysis in the same 6 female subjects who consistently reported their menstrual cycles over the year, and 2 additional subjects who reported them over a six-month period. Sleep onset was later by 9.5 min [7.1–14.1] four days before menstruation onset (p<0.05, d=0.75; Supplementary Fig. S6), and the proportion of REM sleep was higher by 1.0% [0.88–1.34] the day after onset (p<0.05, d=0.75).

Social jetlag (free-day sleep midpoint minus mean work-day sleep midpoint) was 0.88 h [0.65–1.10], ranging from 0.24 to 1.68 h (Supplementary Fig. S7), with 6 of 20 participants having social jetlag ≥1h. Sleep-corrected social jetlag was lower, 0.76 h [0.62–0.90], reduced relative to the raw estimate in 13 of 20 subjects and unchanged in the remaining 7, all of whom had non-positive catch-up sleep. Catch-up sleep was modest, 0.14 h [−0.07–0.37], and was significantly correlated with raw social jetlag (ρ=0.469, p=0.0369).

Sleep regularity was high across participants, SRI=1−SD(sleep midpoint)/12h, with a median of 0.907 [0.873–0.919]. Sex and age effects on inter-individual differences in sleep were assessed via a subject-level regression (metric ∼ sex + age, both z-scored, sex coded female=1/male=0, standardized β and unique R² reported, all 28 tests FDR-corrected together). Effects were concentrated on sleep timing: males had a later sleep onset (β=−1.08, unique R²=0.305, q=0.0345) and later sleep midpoint (β=−1.03, unique R²=0.272, q=0.0345) than females; older age was associated with an earlier sleep offset (β=−0.549, unique R²=0.300, q=0.0345) and earlier sleep midpoint (β=−0.511, unique R²=0.260, q=0.0345).

## Discussion

In this study, we characterized ultralong-term sleep rhythmicity over the course of one year using sqEEG recordings collected in a naturalistic setting. The main implication of these data is that human sleep, when measured repeatedly with EEG in everyday life, is organized across multiple temporal scales (Fig. 5). We found that external cues such as season, weather, and day of week selectively shifted sleep timing and architecture, while evidence for intrinsic multi-day rhythms remains sparse. Thus, long-term EEG monitoring can reveal sleep regulatory patterns that are not accessible from single-night PSG, laboratory protocols, or non-EEG wearable studies enabling only approximation of brain activity in artificial settings.

**Figure 5.**
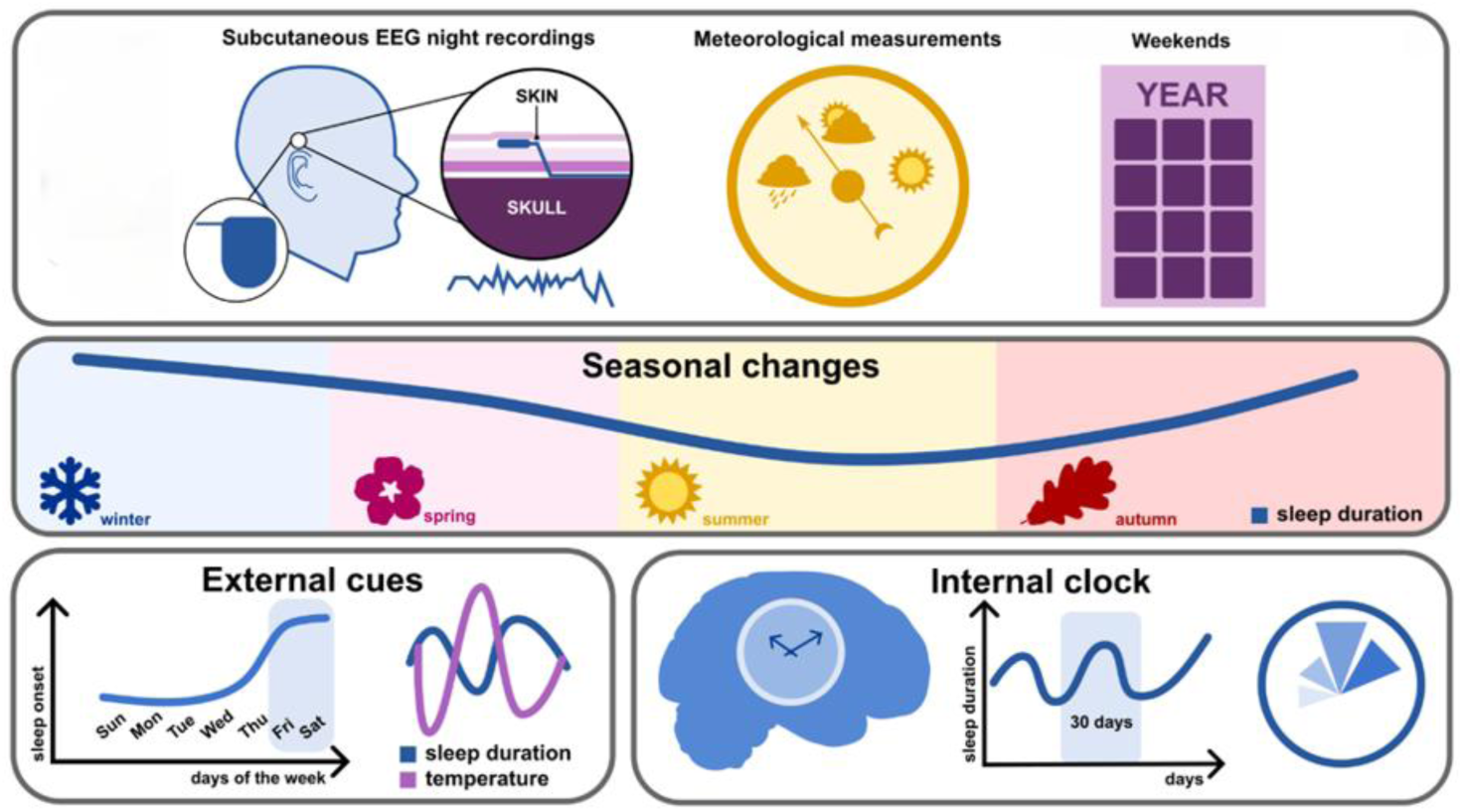
Graphical abstract. Year-long subcutaneous EEG monitoring in 20 healthy adults.

Cycles are a common feature of physiology, influencing everything from hormone release(28) to body temperature(40). Although the circadian clock is well established(41), recent work in other physiological systems suggests slower rhythms operating over weeks or months(40). For sleep research, such rhythms are particularly difficult to study because they require continuous, high-quality recordings collected over long periods in the natural environment. This study builds on previous actigraphy studies that investigated sleep in naturalistic settings(42, 43), including the Whitehall II cohort, which demonstrated the importance of sleep for health and function, as well as the temporal dependency of sleep on multiple factors such as season, light, and temperature(44, 45). Important laboratory studies using melatonin levels also showed the dependency of sleep and circadian biology on environmental factors such as season(46). By adding EEG-derived sleep architecture over a full year, the present work bridges the gap between scalable wearable studies and mechanistically richer sleep recordings.

It is increasingly recognized that human biology is governed not only by 24-hour circadian rhythms but also by ultralong-term multi-day cycles. Research in chronobiology has documented recurring ultralong-period cycles, in the order of days to weeks, across numerous physiological parameters, including heart rate(47), temperature, metabolism(48), and the internal correlations between them(49). The present findings extend this concept to EEG-defined sleep as only direct window into human brain activity. However, the extent to which human sleep, and specifically sleep architecture, is influenced by endogenous multi-day rhythms has been poorly studied, and the relative composition of sleep has rarely been examined, likely due to the limited ability of wearable devices to accurately assess specific stages such as REM sleep(50). We did not find strong evidence of a robust intrinsic cycle in most patients for most sleep macrostructure parameters. However, some subject demonstrated a weak intrinsic cycle between 8 to 60 days for sleep latency and WASO which were not accounted by external cues. This question of long term cycles has often been overlooked and, to date, has primarily been studied in the context of hormonal regulation(28), shift work(51), or sleep restriction(52) for relatively short durations, rather than in naturalistic and ultralong-term settings. Even in the broader literature on animal models, this area remains underexplored, with most studies focusing on genetic mutations or circadian rhythms(41, 53), rather than on multi-day cycles. These findings might have relevant implications for biology and medicine. In conditions affected by sleep, such as dementia(54) and epilepsy(55), recognizing and accounting for multi-day shifts in sleep could improve symptom monitoring and treatment timing. A practical implication is that a single night of sleep, or even several nights sampled over a short window, may not adequately represent a person’s typical sleep architecture or sleep continuity. In clinical research, unmodeled multi-day sleep rhythms could contribute to variability in outcomes such as cognition, mood, seizure occurrence, or response to interventions. Longitudinal sleep tracking could therefore serve as a non-invasive biomarker for forecasting periods of vulnerability or for optimizing the timing of behavioral, pharmacological, or neuromodulatory interventions.

By analyzing a year-long continuous sleep EEG dataset from healthy individuals, we demonstrated that sleep timing and architecture vary systematically across the year. Total sleep duration and REM duration were both reduced in June, while sleep timing shifted seasonally: onset later in July, offset earlier in September, and offset later again in December, alongside increased N3 duration in April and increased REM duration in February. These observations are consistent with previous studies that used non-continuous data to investigate seasonal effects on sleep(56-59). Similar patterns have also been observed with commercially available auxiliary devices, further supporting seasonal modulation of human sleep duration and sleep onset/offset(12, 14, 15), with effects influenced by geographic location(17) and distance from the equator(16). A key contribution of the present study is that seasonal changes in sleep timing can now be interpreted alongside concurrent changes in EEG-defined sleep architecture. The early-summer pattern likely reflects the combined influence of photoperiod, social schedules, temperature, and behavioral adaptation, rather than a single environmental driver.

A key advantage of ultralong-term sleep monitoring in the home environment is the ability to assess the impact of external factors, such as weather and social timing, on sleep patterns, while simultaneously disentangling these effects from potential endogenous modulations of sleep over time. Brighter sunshine was associated with shorter sleep latency, whereas higher temperature was associated with reduced N3 duration. Hot nights were associated with shorter TST and small shifts in N2 proportion. These findings are consistent with prior work showing that environmental temperature and light exposure can shape sleep timing and sleep continuity(12, 60), but extend this literature by showing that environmental influences are reflected not only in sleep timing and continuity, but also in EEG-defined sleep architecture.

Our findings regarding later sleep onset and offset during weekends are consistent with previous research(15, 61-63). However, few studies have examined this effect over an extended period in a natural environment using EEG (full description in Table S1). The weekend shift represented one of the strongest external influences on sleep timing, with sleep onset, sleep offset, and sleep midpoint delayed by approximately one hour on Friday and Saturday nights relative to Monday–Thursday nights. This pattern is clinically and behaviorally relevant because repeated weekly shifts in sleep timing may contribute to circadian misalignment, social jetlag, and variability in daytime functioning. These findings show that sleep timing shifted consistently later on weekend nights, with onset, offset, and sleep midpoint all delayed on Friday and Saturday. This shift was accompanied by a significant change in sleep latency, indicating that later weekend sleep is linked to sleep continuity. Together, these findings leave open whether the weekend delay in timing reflects reduced social constraints or intrinsic sleep preferences.

To address the challenge of multiple covarying influences in a real-world dataset, we performed a PCA to identify the dominant dimensions of nightly sleep variability. This analysis showed that sleep continuity and architecture loaded together on PC1, while sleep timing formed a largely separate axis on PC2, indicating that these are partly independent dimensions of variation. Projecting each longitudinal driver onto these components (a descriptive analysis without a significance test) showed that hot nights had the largest projected share on every one of the first five components, followed by the seasonal trend and then the day-of-week effect, with a broadly similar proportional split across components rather than concentration on any single one. The intrinsic-cycle driver’s projected share was negligible by comparison on every component; because its magnitude is a phase-locking value rather than a raw deviation in the metrics’ native units, this near-zero share reflects a scale mismatch rather than an absence of structural alignment with these components. PCA therefore provides a useful framework for integrating the effects of multiple interacting factors and for interpreting sleep as a multidimensional system rather than as a collection of independent variables. We also explored the influence of the menstrual cycle on sleep. As only a small number of participants provided menstrual cycle data for at least six months, this analysis should be considered exploratory. We observed that sleep onset was delayed in the days preceding menstruation onset, and that REM sleep proportion increased on the day after onset, indicating cyclical modulation of both sleep timing and sleep architecture across the menstrual cycle. These observations are consistent with previous evidence that reproductive physiology can influence sleep architecture (28). However, the small sample size and restricted age range of the subgroup substantially limit generalizability. These findings should therefore be regarded as hypothesis-generating rather than definitive evidence for menstrual-cycle effects on long-term EEG-derived sleep dynamics.

The social jetlag and sleep-regularity analyses further emphasize that real-world sleep is shaped by both intrinsic biological factors and social schedules. Although participants generally exhibited high sleep regularity, 6 of 20 participants (30%) demonstrated social jetlag of at least one hour. This finding suggests that even in a relatively healthy cohort, social obligations impose a measurable weekly displacement of sleep timing in a meaningful subset of individuals. Integrating social-timing measures with EEG-derived sleep architecture may help disentangle the effects of behavioural schedules from those of intrinsic physiological rhythms in future studies.

Several limitations should be considered. First, the cohort comprised only 20 participants recruited within a single country. Although the study generated a very dense longitudinal dataset comprising over 7000 recorded nights, replication in larger and more geographically diverse cohorts will be necessary to confirm the observed year-long trends and multi-day rhythmic patterns. Because the study was conducted in Denmark, where seasonal changes in photoperiod and weather differ substantially from many other regions, the magnitude and timing of the observed seasonal effects may not generalize directly to other populations. In addition, some participants travelled outside Denmark for a total of 38 confirmed days during the study period. Although this represents only a small fraction of the total observation time, travel-related changes in environment, daylight exposure, and behaviour may have contributed to individual variability. Second, the menstrual-cycle analysis was based on a small subgroup of female participants aged 21–40 years and should therefore be regarded as exploratory. Similarly, the age- and sex-related analyses were not powered to support strong subgroup inferences and should be interpreted cautiously. Third, the observational design precludes causal inference regarding the relationships among environmental, social, and endogenous influences on sleep architecture, and residual confounding remains possible despite adjustment for major external factors. Finally, sqEEG differs from full polysomnography in electrode coverage and available physiological channels, and sleep-stage estimates remain dependent on the performance of the sleep-scoring pipeline.

These limitations should be considered alongside several important strengths. We characterized EEG-derived sleep architecture continuously over an entire year in a naturalistic setting. The study achieved exceptionally high recording compliance and generated an unprecedented longitudinal dataset comprising more than 7,000 nights of EEG recordings. The use of EEG-based sleep staging, rather than surrogate sleep measures, enabled direct assessment of sleep architecture over timescales ranging from days to seasons. Missing nights were excluded from the relevant analyses, and no outlier exclusion procedures were applied. This preserves the naturalistic nature of the dataset but also means that extreme observations, whether arising from unusual behaviours, environmental exposures, or technical factors, contributed to the reported estimates. The dense within-subject sampling provided sufficient temporal resolution to identify multi-day rhythmic patterns that would be difficult to detect in conventional short-duration studies. Finally, the simultaneous evaluation of intrinsic rhythms, environmental influences, and social timing offers a uniquely comprehensive perspective on the temporal organization of human sleep under real-world conditions.

In conclusion, year-long sqEEG monitoring reveals that human sleep in everyday life is shaped by external environmental influences, social cues and intrinsic factors. The broader significance of this work is that sleep should be understood as a longitudinal physiological process rather than a static nightly state. Recognizing these temporal structures may improve the design of sleep studies, reduce unexplained variability in clinical and cognitive outcomes, and support future rhythm-aware approaches to monitoring and intervention.

Beyond healthy sleep physiology, these findings may have important implications for neurological disorders in which sleep plays a central role. In epilepsy, seizure risk fluctuates with sleep stage and circadian timing, and multi-day rhythms in brain activity have been shown to predict periods of increased seizure likelihood. Long-term tracking of sleep architecture may therefore provide a non-invasive means of identifying periods of heightened vulnerability and optimizing the timing of interventions. In dementia, disruption of slow-wave and REM sleep has been linked to impaired clearance of amyloid and tau proteins, raising the possibility that longitudinal sleep monitoring could serve as an early biomarker of disease-relevant change. Similar opportunities may exist in mood disorders, chronic pain, and other conditions in which altered sleep regulation contributes to symptom burden and disease progression.

Future studies should investigate how wearable-derived circadian and behavioral metrics relate to continuously recorded EEG sleep architecture. Integrating EEG, actigraphy, light exposure, temperature, hormonal markers, and subjects-reported outcomes may help disentangle endogenous rhythms from environmental and behavioral influences and provide a more complete understanding of long-term sleep regulation. Such multimodal datasets could also clarify the mechanisms underlying the intrinsic multi-day sleep cycles observed in this study.

Finally, an important next step will be to determine whether these ultralong-term changes are preserved, altered, or disrupted in neurological and psychiatric disorders. More broadly, the ability to monitor sleep continuously over months to years opens new opportunities for precision sleep medicine, in which interventions can be adapted not only to the individual, but also to their dynamic physiological state over time.

## Contributors

Tamir Avigdor (Conceptualization: Equal; Formal analysis: Lead; Investigation: Equal; Methodology: Lead; Software: Lead; Validation: Equal; Visualization: Lead; Writing – original draft: Lead; Writing – review & editing: Equal), Jonas Duun-Henriksen (Conceptualization: Supporting; Data curation: Equal; Investigation: Supporting; Validation: Supporting; Writing – review & editing: Supporting), Esben Ahrens (Data curation: Supporting), Alyssa Ho (Data curation: Supporting; Formal analysis: Supporting; Methodology: Supporting; Visualization: Supporting; Writing – review & editing: Supporting), Matthew Moye (Data curation: Supporting; Visualization: Supporting; Writing – review & editing: Supporting), David Carlson (Investigation: Supporting; Methodology: Supporting), Birgit Frauscher (Conceptualization: Equal; Formal analysis: Supporting; Investigation: Equal; Methodology: Equal; Supervision: Equal; Validation: Equal; Writing – original draft: Equal; Writing – review & editing: Equal), Sandor Beniczky (Conceptualization: Lead; Investigation: Lead; Methodology: Supporting; Project administration: Lead; Supervision: Lead; Validation: Supporting; Writing – original draft: Lead; Writing – review & editing: Lead).

T.A. & S.B. accessed and verified the underlying data.

All authors read and approved the final version of the manuscript.

## Data sharing statement

Data belongs to UNEEG and cannot be shared. The code supporting the analysis and findings of this study is available from the authors upon request.

## Acknowledgements

Thanks to funding sources for our study. Filadelfia Research Fund (Grant F2024.04) was awarded to Sándor Beniczky. Duke University startup fund & Canadian Institutes of Health Research (PJT-175056) was awarded to Birgit Frauscher.

## References

1. Richards J, Gumz ML. Mechanism of the circadian clock in physiology. Am J Physiol Regul Integr Comp Physiol. 2013;304(12):R1053–64.

2. Pelayo R, Hodgson N, Guilleminault C. Chapter 34: the history of sleep medicine. Handb Clin Neurol. 2010;95:547–56.

3. Pelayo R, Guilleminault C. History of sleep research. The neuroscience of sleep. 2009;3.

4. Davis H, Davis PA, Loomis AL, Harvey EN, Hobart G. Changes in Human Brain Potentials during the Onset of Sleep. Science. 1937;86(2237):448–50.

5. Pantev C. Evoked and induced gamma-band activity of the human cortex. Brain Topogr. 1995;7(4):321–30.

6. Academy of Sleep Medicine A. The AASM Manual for the Scoring of Sleep and Associated Events Summary of Updates in Version 2.52020.

7. Ghegan MD, Angelos PC, Stonebraker AC, Gillespie MB. Laboratory versus portable sleep studies: a meta-analysis. Laryngoscope. 2006;116(6):859–64.

8. Newell J, Mairesse O, Verbanck P, Neu D. Is a one-night stay in the lab really enough to conclude? First-night effect and night-to-night variability in polysomnographic recordings among different clinical population samples. Psychiatry Res. 2012;200(2-3):795–801.

9. Levendowski DJ, Zack N, Rao S, Wong K, Gendreau M, Kranzler J, et al. Assessment of the test-retest reliability of laboratory polysomnography. Sleep Breath. 2009;13(2):163–7.

10. Kleitman N. Sleep and wakefulness as alternating phases in the cycle of existence. Chicago,: Univ. of Chicago Press; 1939. xii, 638 p. p.

11. de Zambotti M, Cellini N, Goldstone A, Colrain IM, Baker FC. Wearable Sleep Technology in Clinical and Research Settings. Med Sci Sports Exerc. 2019;51(7):1538–57.

12. Mattingly SM, Grover T, Martinez GJ, Aledavood T, Robles-Granda P, Nies K, et al. The effects of seasons and weather on sleep patterns measured through longitudinal multimodal sensing. NPJ Digit Med. 2021;4(1):76.

13. Li L, Nakamura T, Hayano J, Yamamoto Y. Seasonal Sleep Variations and Their Association With Meteorological Factors: A Japanese Population Study Using Large-Scale Body Acceleration Data. Front Digit Health. 2021;3:677043.

14. Luong N, Mark G, Kulshrestha J, Aledavood T. Sleep During the COVID-19 Pandemic: Longitudinal Observational Study Combining Multisensor Data With Questionnaires. JMIR Mhealth Uhealth. 2024;12:e53389.

15. Oskarsdottir M, Islind AS, August E, Arnardottir ES, Patou F, Maier AM. Importance of Getting Enough Sleep and Daily Activity Data to Assess Variability: Longitudinal Observational Study. JMIR Form Res. 2022;6(2):e31807.

16. Scott H, Lechat B, Sansom K, Pinilla L, Manners J, Phillips AJK, et al. Variations in sleep duration and timing: Weekday and seasonal variations in sleep are common in an analysis of 73 million nights from an objective sleep tracker. Sleep. 2025.

17. Willoughby AR, Alikhani I, Karsikas M, Chua XY, Chee MWL. Country differences in nocturnal sleep variability: Observations from a large-scale, long-term sleep wearable study. Sleep Med. 2023;110:155–65.

18. Yetish G, Kaplan H, Gurven M, Wood B, Pontzer H, Manger PR, et al. Natural sleep and its seasonal variations in three pre-industrial societies. Curr Biol. 2015;25(21):2862–8.

19. Koch M, Matzke I, Huhn S, Sie A, Boudo V, Compaore G, et al. Assessing the Effect of Extreme Weather on Population Health Using Consumer-Grade Wearables in Rural Burkina Faso: Observational Panel Study. JMIR Mhealth Uhealth. 2023;11:e46980.

20. Basner M, Dinges DF, Mollicone D, Ecker A, Jones CW, Hyder EC, et al. Mars 520-d mission simulation reveals protracted crew hypokinesis and alterations of sleep duration and timing. Proc Natl Acad Sci U S A. 2013;110(7):2635–40.

21. Flynn-Evans EE, Barger LK, Kubey AA, Sullivan JP, Czeisler CA. Circadian misalignment affects sleep and medication use before and during spaceflight. NPJ Microgravity. 2016;2:15019.

22. Blunden S, Magee C, Clarkson L, Searle A, Banks S, Olds T. Interindividual and intraindividual variability in adolescent sleep patterns across an entire school term: A pilot study. Sleep Health. 2019;5(6):546–54.

23. Steinach M, Kohlberg E, Maggioni MA, Mendt S, Opatz O, Stahn A, et al. Sleep Quality Changes during Overwintering at the German Antarctic Stations Neumayer II and III: The Gender Factor. PLoS One. 2016;11(2):e0150099.

24. Lee T, Cho Y, Cha KS, Jung J, Cho J, Kim H, et al. Accuracy of 11 Wearable, Nearable, and Airable Consumer Sleep Trackers: Prospective Multicenter Validation Study. JMIR Mhealth Uhealth. 2023;11:e50983.

25. Birrer V, Elgendi M, Lambercy O, Menon C. Evaluating reliability in wearable devices for sleep staging. NPJ Digit Med. 2024;7(1):74.

26. Robbins R, Weaver MD, Sullivan JP, Quan SF, Gilmore K, Shaw S, et al. Accuracy of Three Commercial Wearable Devices for Sleep Tracking in Healthy Adults. Sensors (Basel). 2024;24(20).

27. Helge AW, Arguissain FG, Lechner L, Gritsch G, Duun-Henriksen J, Ahrens E, et al. Longitudinal, EEG-based assessment of sleep in people with epilepsy: An automated sleep staging algorithm non-inferior to human raters. Clin Neurophysiol Pract. 2025;10:30–9.

28. Alzueta E, Baker FC. The Menstrual Cycle and Sleep. Sleep Med Clin. 2023;18(4):399–413.

29. Duun-Henriksen J, Baud M, Richardson MP, Cook M, Kouvas G, Heasman JM, et al. A new era in electroencephalographic monitoring? Subscalp devices for ultra-long-term recordings. Epilepsia. 2020;61(9):1805–17.

30. Ahrens E, Jennum P, Duun-Henriksen J, Borregaard HWS, Nielsen SS, Taptiklis N, et al. The Ultra-Long-Term Sleep study: Design, rationale, data stability and user perspective. J Sleep Res. 2024;33(6):e14197.

31. Perslev M, Darkner S, Kempfner L, Nikolic M, Jennum PJ, Igel C. U-Sleep: resilient high-frequency sleep staging. NPJ Digit Med. 2021;4(1):72.

32. Wittmann M, Dinich J, Merrow M, Roenneberg T. Social jetlag: misalignment of biological and social time. Chronobiol Int. 2006;23(1-2):497–509.

33. Jankowski KS. Social jet lag: Sleep-corrected formula. Chronobiol Int. 2017;34(4):531–5.

34. Wichmann J, Andersen Z, Ketzel M, Ellermann T, Loft S. Apparent temperature and cause-specific emergency hospital admissions in Greater Copenhagen, Denmark. PLoS One. 2011;6(7):e22904.

35. Karoly PJ, Rao VR, Gregg NM, Worrell GA, Bernard C, Cook MJ, et al. Cycles in epilepsy. Nat Rev Neurol. 2021;17(5):267–84.

36. Baud MO, Kleen JK, Mirro EA, Andrechak JC, King-Stephens D, Chang EF, et al. Multi-day rhythms modulate seizure risk in epilepsy. Nat Commun. 2018;9(1):88.

37. Xiong W, Stirling RE, Payne DE, Nurse ES, Kameneva T, Cook MJ, et al. Forecasting seizure likelihood from cycles of self-reported events and heart rate: a prospective pilot study. EBioMedicine. 2023;93:104656.

38. Maris E, Oostenveld R. Nonparametric statistical testing of EEG- and MEG-data. J Neurosci Methods. 2007;164(1):177–90.

39. Rana KD, Vaina LM, Hamalainen MS. A fast statistical significance test for baseline correction and comparative analysis in phase locking. Front Neuroinform. 2013;7:3.

40. Harding C, Pompei F, Bordonaro SF, McGillicuddy DC, Burmistrov D, Sanchez LD. The daily, weekly, and seasonal cycles of body temperature analyzed at large scale. Chronobiology International. 2019;36(12):1646–57.

41. Sanchez REA, Kalume F, de la Iglesia HO. Sleep timing and the circadian clock in mammals: Past, present and the road ahead. Semin Cell Dev Biol. 2022;126:3–14.

42. Ferrie JE, Shipley MJ, Cappuccio FP, Brunner E, Miller MA, Kumari M, et al. A prospective study of change in sleep duration: Associations with mortality in the Whitehall II Cohort. Sleep. 2007;30(12):1659–66.

43. Ferrie JE, Shipley MJ, Akbaraly TN, Marmot MG, Kivimäki M, Singh-Manoux A. Change in Sleep Duration and Cognitive Function: Findings from the Whitehall II Study. Sleep. 2011;34(5):565–73a.

44. Juda M, Vetter C, Roenneberg T. The Munich ChronoType Questionnaire for Shift-Workers (MCTQShift). J Biol Rhythm. 2013;28(2):130–40.

45. Okamoto-Mizuno K, Mizuno K. Effects of thermal environment on sleep and circadian rhythm. J Physiol Anthropol. 2012;31.

46. Wehr TA, Moul DE, Barbato G, Giesen HA, Seidel JA, Barker C, et al. Conservation of Photoperiod-Responsive Mechanisms in Humans. American Journal of Physiology. 1993;265(4):R846–R57.

47. Dijk DJ, Duffy JF. Novel Approaches for Assessing Circadian Rhythmicity in Humans: A Review. J Biol Rhythms. 2020;35(5):421–38.

48. Refinetti R. Circadian rhythmicity of body temperature and metabolism. Temperature (Austin). 2020;7(4):321–62.

49. Thornton C, Smith BC, Besne GM, Wang Y, editors. Using Wearable Sensors to Capture the Synchrony of Circadian Rhythms Across Physiological Processes in Free-living Conditions. 2024 IEEE BioSensors Conference (BioSensors); 2024: IEEE.

50. Danzig R, Wang M, Shah A, Trotti LM. The wrist is not the brain: Estimation of sleep by clinical and consumer wearable actigraphy devices is impacted by multiple patient- and device-specific factors. J Sleep Res. 2020;29(1):e12926.

51. Wu QJ, Sun H, Wen ZY, Zhang M, Wang HY, He XH, et al. Shift work and health outcomes: an umbrella review of systematic reviews and meta-analyses of epidemiological studies. J Clin Sleep Med. 2022;18(2):653–62.

52. Banks S, Dinges DF. Behavioral and physiological consequences of sleep restriction. J Clin Sleep Med. 2007;3(5):519–28.

53. Bloch G, Barnes BM, Gerkema MP, Helm B. Animal activity around the clock with no overt circadian rhythms: patterns, mechanisms and adaptive value. Proc Biol Sci. 2013;280(1765):20130019.

54. Xu W, Tan CC, Zou JJ, Cao XP, Tan L. Sleep problems and risk of all-cause cognitive decline or dementia: an updated systematic review and meta-analysis. J Neurol Neurosurg Psychiatry. 2020;91(3):236–44.

55. Sheybani L, Frauscher B, Bernard C, Walker MC. Mechanistic insights into the interaction between epilepsy and sleep. Nat Rev Neurol. 2025;21(4):177–92.

56. Al Lawati I, Zadjali F, Al-Abri MA. Seasonal variation and sleep patterns in a hot climate Arab Region. Sleep Breath. 2023;27(1):355–62.

57. Evans DS, Snitker S, Wu SH, Mody A, Njajou OT, Perlis ML, et al. Habitual sleep/wake patterns in the Old Order Amish: heritability and association with non-genetic factors. Sleep. 2011;34(5):661–9.

58. Honma K, Honma S, Kohsaka M, Fukuda N. Seasonal variation in the human circadian rhythm: dissociation between sleep and temperature rhythm. Am J Physiol. 1992;262(5 Pt 2):R885-91.

59. Kume Y, Makabe S, Singha-Dong N, Vajamun P, Apikomonkon H, Griffiths J. Seasonal effects on the sleep-wake cycle, the rest-activity rhythm and quality of life for Japanese and Thai older people. Chronobiol Int. 2017;34(10):1377–87.

60. Chevance G, Minor K, Vielma C, Campi E, O’Callaghan-Gordo C, Basagana X, et al. A systematic review of ambient heat and sleep in a warming climate. Sleep Med Rev. 2024;75:101915.

61. Crowley SJ, Carskadon MA. Modifications to weekend recovery sleep delay circadian phase in older adolescents. Chronobiol Int. 2010;27(7):1469–92.

62. Misiunaite I, Eastman CI, Crowley SJ. Circadian Phase Advances in Response to Weekend Morning Light in Adolescents With Short Sleep and Late Bedtimes on School Nights. Front Neurosci. 2020;14:99.

63. Paine SJ, Gander PH. Differences in circadian phase and weekday/weekend sleep patterns in a sample of middle-aged morning types and evening types. Chronobiol Int. 2016;33(8):1009–17.

