## Supporting information for "Beyond circadian: a yearlong prospective cohort study into human sleep using subcutaneously implanted electroencephalography"

### Supporting Materials

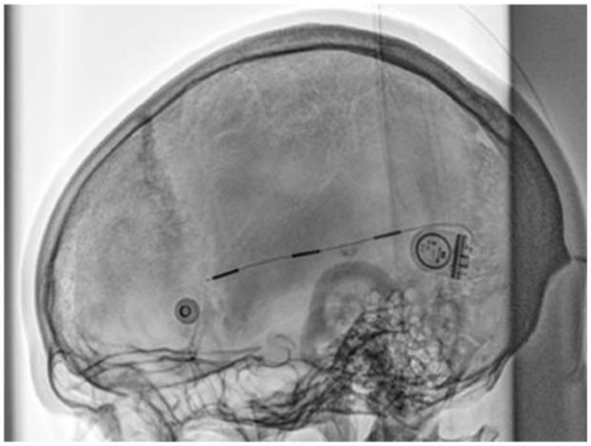

**Figure S1.** Subcutaneous electroencephalogram Post-implantation X-ray. Showing the device positioned over the superior temporal region. The implant lead is placed deep to the galea and directed posteriorly to cover the temporal lobe, while the housing is positioned subcutaneously and bent approximately 90° to help pull the lead further forward. Although different lead trajectories can be used to target various cortical regions, the device contains only a single lead.

Djurhuus BD, Viana PF, Ahrens E, et al. Minimally invasive surgery for placement of a subcutaneous EEG implant. Front Surg. 2023 Nov 9;10:1304343. doi: 10.3389/fsurg.2023.1304343 - published under the Creative Commons Attribution License (CC BY 4.0).

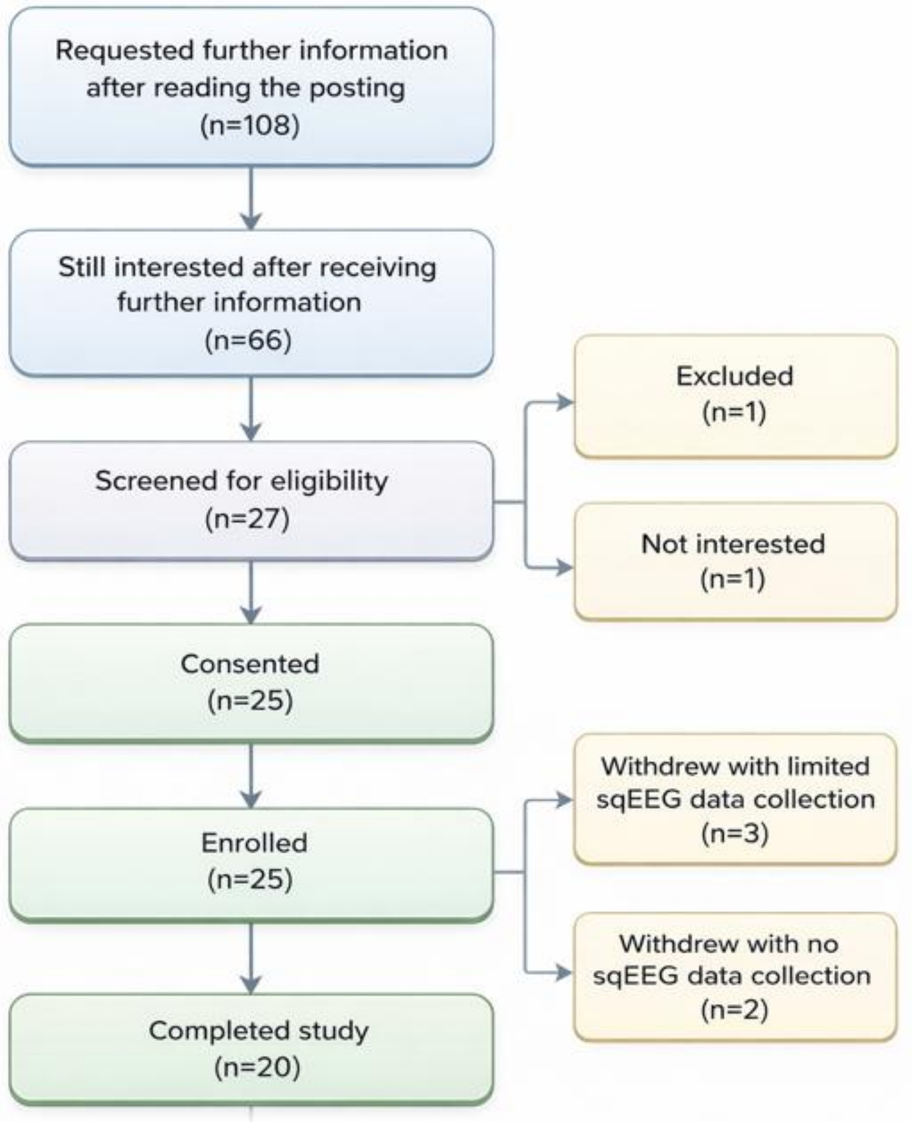


**Figure S2.** Participant selection flowchart adapted from the original study (Ahrens et al., 2024).

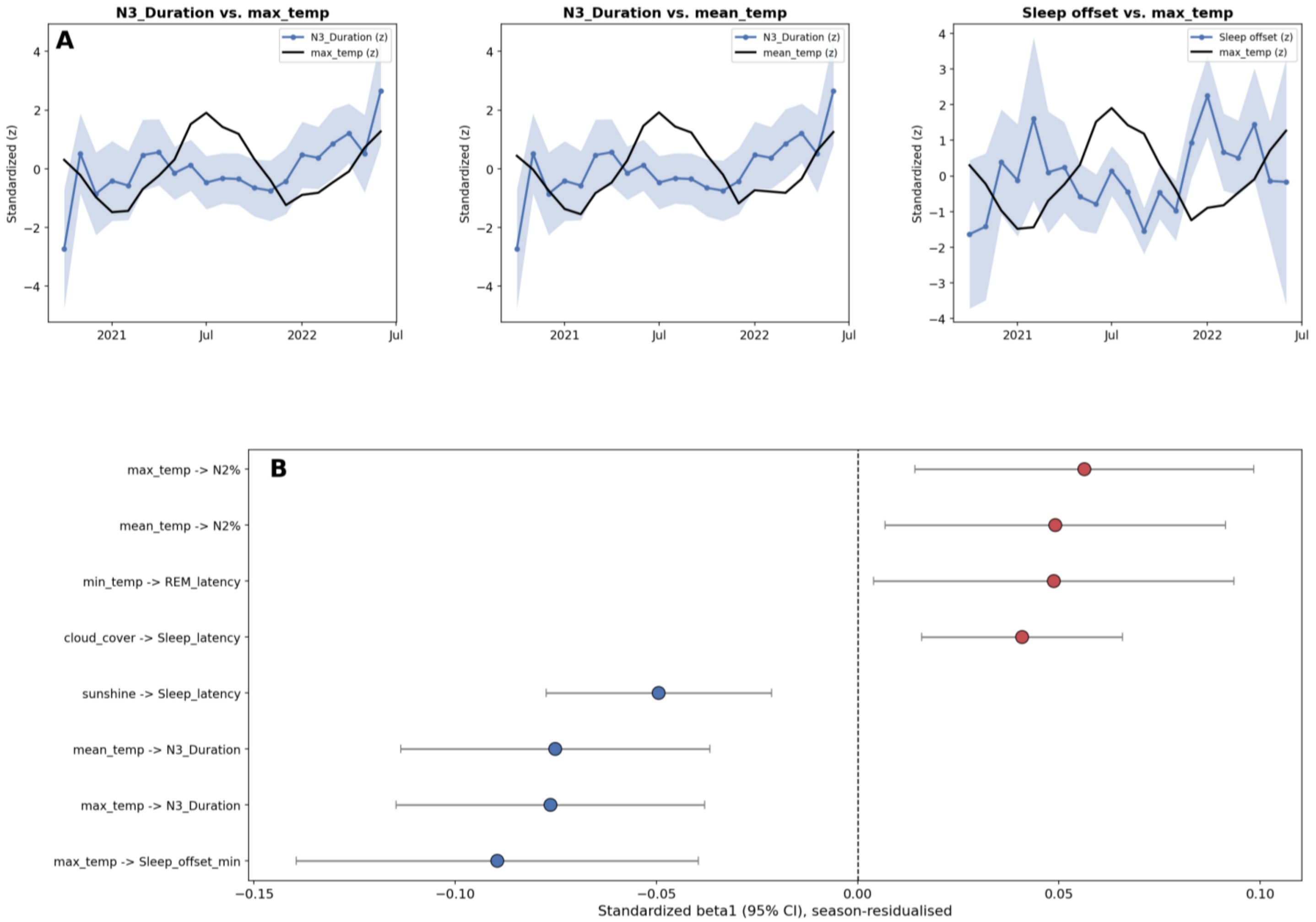


**Figure S3. Weather and sleep.** (A) The three example sleep metric–weather parameter, both series are z-scored to their own mean and SD and plotted on a single shared axis (standardized units). The blue line and shaded band are the cohort mean ± SEM of the sleep metric, aggregated by calendar month, the black line is the mean weather parameter for that same calendar month, restricted to the months the sleep-metric series covers. (B) Standardized fixed-effect beta coefficients (± 95% CI) from linear mixed-effects models (sleep metric ~ weather parameter + month + (1|subject)) relating each weather parameter to each sleep metric, fit jointly across all participants with subject as a random intercept. Benjamini-Hochberg false discovery rate (FDR) correction applied jointly across all pairs tested in this analysis (α=0.05).

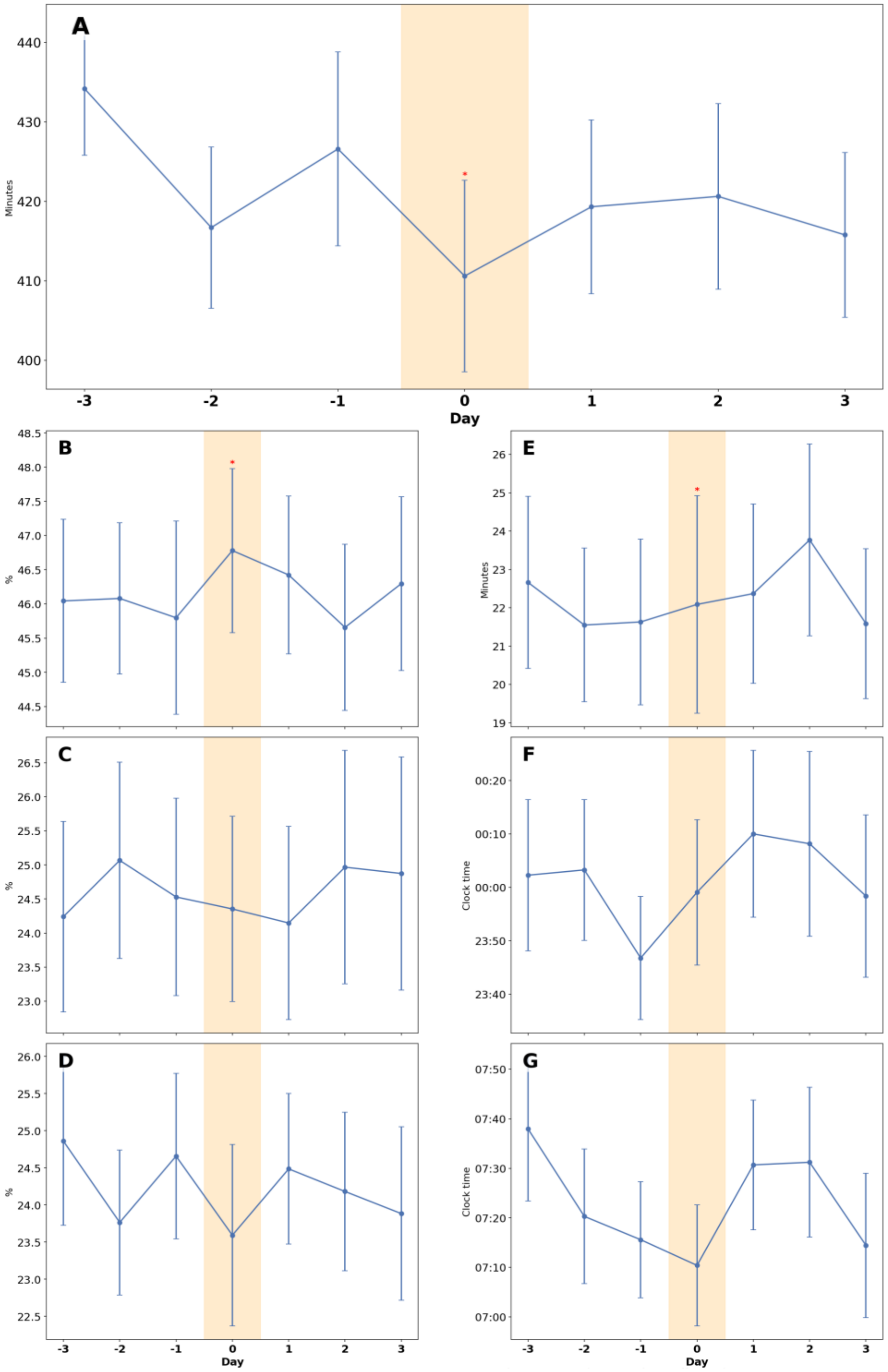

**Figure S4. Impact of excessively hot nights on sleep architecture.** Sleep metrics were compared between nights with heat stress, defined as nighttime temperature >20°C, and each participant's sleep during the preceding three nights. Green bars indicate sleep values on the heat-stress night. Statistical significance was assessed using a paired permutation test comparing the heat-stress night with the participant-specific preceding 3-day distribution. Significant differences are indicated by *p<0.05, **p<0.01, and ***p<0.001. Abbreviations: WASO, wake after sleep onset; R, rapid eye movement sleep.

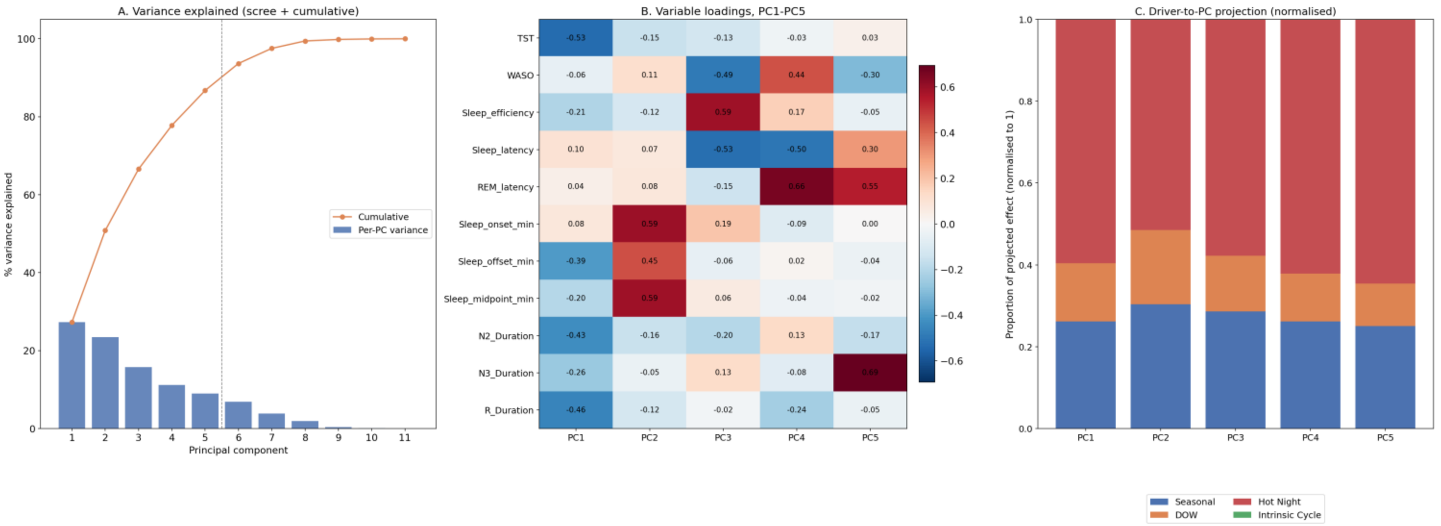


**Figure S5. Principel component analysis** (A) Scree plot of PCA on within-subject z-scored sleep variables; bars show variance explained by each PC and the line shows cumulative variance (first 5 PCs summarize the dominant latent structure).(B) Heatmap of variable on PC1–PC5 (red/blue = positive/negative contribution), with numeric coefficients to identify which sleep features define each latent axis.(C) Projection of analysis-source effect vectors (Seasonal, day of three weeks (DOW), Hot Night, Intrinsic Cycle) onto the PCA basis, computed as Σ|effect × loading| across variables for each PC; stacked bars show how strongly each source maps onto each latent component.

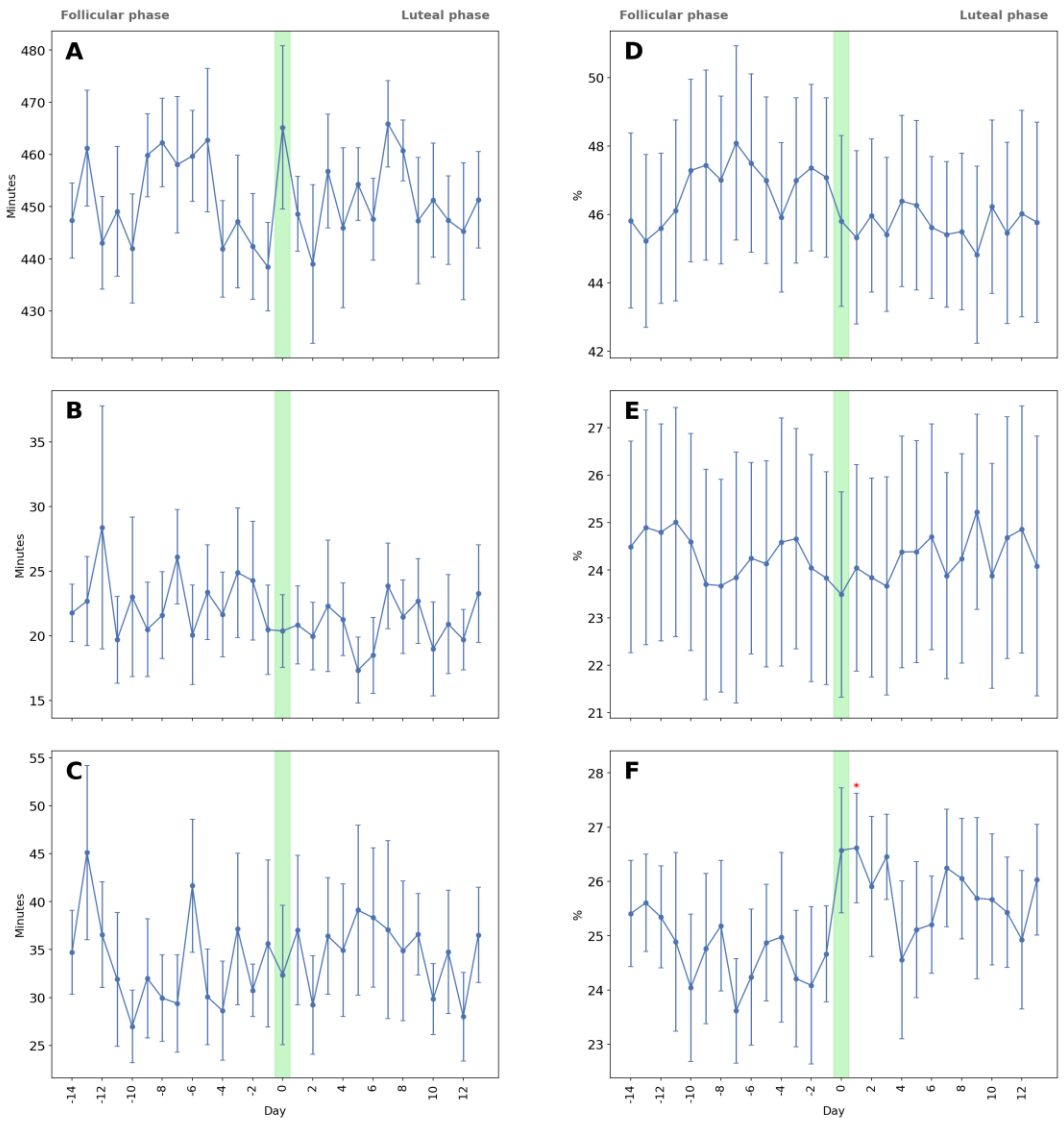


**Figure S6. Sleep parameters aligned to the start of menstruation.** Sleep parameters are shown aligned to menstruation onset, marked as day zero (the 28-day window spans day −14 to day +13): (A) total sleep time (TST); (B) wake after sleep onset (WASO); (C) sleep latency; (D) N2 sleep percentage; (E) N3 sleep percentage; (F) REM sleep percentage. Plots display the cohort mean ±

SEM of the raw values across subjects. A paired, two-tailed (p<0.05), cluster-based sign-flip permutation test (10,000 iterations) was performed on each subjects.. Green marks the starting day of menstruation, as manually noted into the auxiliary device.

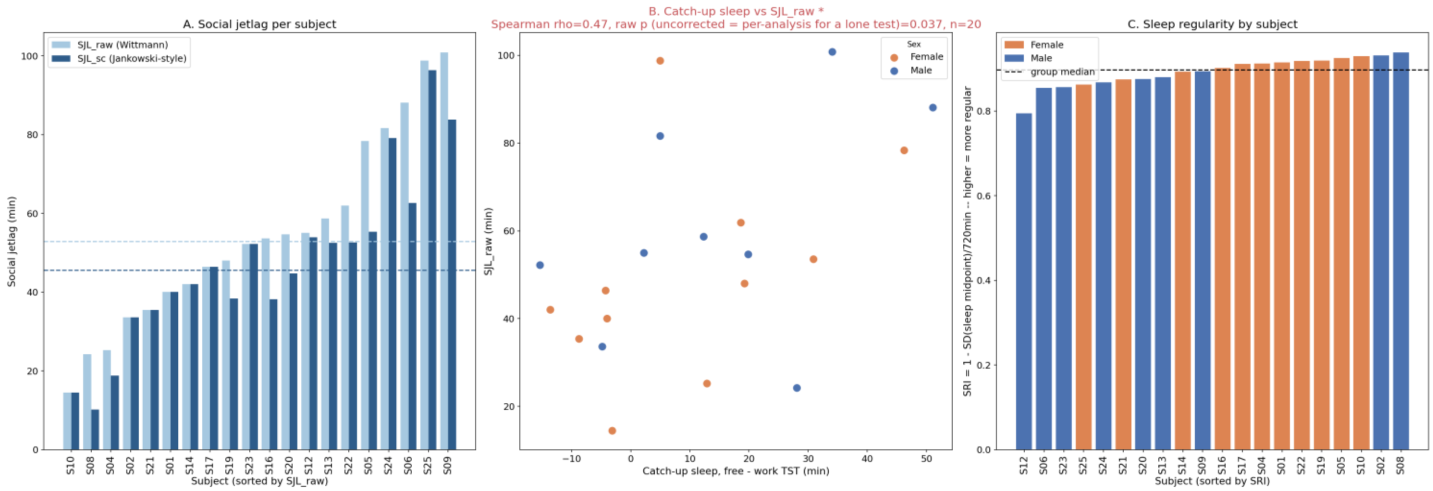


**Figure S7. Social jetlag and sleep regularity across participants.** A. Between-subject variability in raw social jetlag (SJL_raw) and sleep-corrected social jetlag (SJL_sc). Dashed lines indicate the cohort median for each measure. B. Association between catch-up sleep and SJL_raw, assessed using Spearman rank correlation. Full statistics are reported in the text. C. Sleep Regularity Index (SRI) for each participant, sorted by value. The dashed line indicates the cohort median SRI.

**Table. S1. Literature review.** provides a structured overview of peer-reviewed studies that have tracked sleep for > 3 weeks. Each row corresponds to a single study, while the columns are organized to let readers quickly compare methodological choices, sample characteristics, and key outcomes.

| Title | Year | Center | Sample Size | Age | Method | Study Duration |
|---|---|---|---|---|---|---|
| Comparison between subjective and actigraphic measurement of sleep and sleep rhythms | 2002 | University of Surrey, Guildford, UK | 49 | 46.6±12 .2 years | Actigraphy and daily questionnaires | 4 weeks |
| Changes in sleep–wake cycle during the period from late pregnancy to puerperium identified through the wrist actigraph and sleep logs | 2002 | Kyushu University School of Health Sciences, Fukuoka-shi, Japan | 4 | | Actigraphy and daily sleep logs | 34 weeks of gestation to 3 months after delivery |
| Longitudinal study for sleep–wake behaviours of mothers from pre-partum to post-partum using actigraph and sleep logs | 2002 | University of East Asia, Yamaguchi, Japan | 10 | 29.5 ± 2.2 years | Actigraphy and daily diary | 20 weeks (5th week before delivery to 15 weeks after delivery) |
| Sleep quality in professional ballet dancers | 2009 | Charité-Universitätsmedizin Berlin, Berlin, Germany | 24 | 27 ± 5 years | Actigraphy and daily sleep logs | 67 days |
| Working Memory Capacity is Decreased in Sleep-Deprived Internal Medicine Residents | 2009 | University of Minnesota and Regions Hospital, St. Paul, MN, USA | 39 | 28.5 ± 3.97 years | Actigraphy | 2 months |
| Longitudinal Study of Sleep Patterns of United States Military Academy Cadets | 2010 | Naval Postgraduate School, Monterey, CA, USA | 55-75 (depending on session) | | Actigraphy | 4x30 days during fall semester, and 4x30 days during spring semester |
| Marital/Cohabitation Status and History in Relation to Sleep in Midlife Women | 2010 | University of Pittsburgh, Pittsburgh, PA, USA | 367 | | Actigraphy | Entire menstrual cycle or 35 days |
| Normative longitudinal maternal sleep: the first four postpartum months | 2010 | West Virginia University, Morgantown, West Virginia, USA | 66 | 27.3 ± 5.8 years | Actigraphy | 8-12 weeks |
| Predicting sleep quality from stress and prior sleep – A study of day-to-day covariation across sixweeks | 2012 | Karolinska Institutet, Stockholm, Sweden | 50 | 18-61 years | Actigraphy and Karolinska Sleep Questionnaire | 6 weeks |

| | | | | | | |
|---|---|---|---|---|---|---|
| Mars 520-d mission simulation reveals protracted crew hypokinesis and alterations of sleep duration and timing | 2013 | State Scientific Center of the Russian Federation–IBMP of the Russian Academy of Sciences (RAS) | 6 | | Actigraphy | 520 days |
| Sleep Disturbance and Neurobehavioral Performance among Postpartum Women | 2013 | West Virginia University, Morgantown, WV, USA | 70 postpartum, 9 nulliparous | 22.2-34.8 years | Actigraphy | 12 weeks |
| Actigraphy-assessed sleep during school and vacation periods: a naturalistic study of restricted and extended sleep opportunities in adolescents | 2013 | The University of Melbourne, Melbourne, Australia | 146 | 16.2 ± 1.0 years | Actigraphy | 4 weeks |
| Sleep duration and weight change in midlife women: The SWAN Sleep Study | 2013 | Chicago, southwest Michigan, Pittsburgh, Oakland (CA) | 310 | 49.7 ± 2.0 years | Actigraphy + 3 nights PSG | Entire menstrual cycle or 35 days |
| Actigraphic and self-reported sleep quality in women: associations with ovarian hormones and mood | 2015 | Sunnybrook Health Sciences Center, Toronto, ON, Canada | 13 | 34 ± 5.7 years | Actigraphy | 42 days |
| A Cognitive Vulnerability Model of Sleep and Mood in Adolescents under Naturalistically Restricted and Extended Sleep Opportunities | 2015 | The University of Melbourne, Melbourne, Australia | 146 | 16.2 ± 1.0 years | Actigraphy | 4 weeks |
| Prevalence of Sleep Deficiency and Hypnotic Use Among Astronauts Before, During and After Spaceflight: An Observational Study | 2015 | Harvard Medical School, Boston, MA, USA | 75 | 46.4 ± 4.5 years | Actigraphy | |
| Fighting fire and fatigue: sleep quantity and quality during multi-day wildfire suppression | 2015 | Deakin University Burwood, VIC, Australia | 41 | 39.4 ± 12.5 years | Actigraphy | 4 weeks |
| Circadian misalignment affects sleep and medication use before and during spaceflight | 2016 | NASA Ames Research Center, Moffett Field, CA, USA | 21 | 46.7 ± 3.9 years | Actigraphy | 59–214 days |
| Cognitive consequences of sleep deprivation, shiftwork, and heat exposure for underground miners | 2016 | Laurentian University, Sudbury, ON, Canada | 19 | 41.5 ± 5.1 years | Actigraphy, sleep questionnaires (KSS) | One rotating shift (~30 days) |
| The Impact of Wearable Device Enabled Health Initiative on Physical Activity and Sleep | 2016 | Indian Institute of Technology, New Delhi, India | 506 | 23-67 years | Actigraphy | 1 year |
| The longer the better: Sleep–wake patternsduring preparation of the World Rowing Junior Championships | 2016 | Ruhr University Bochum, Faculty of Sport Science, Bochum, Germany | 18 with objective sleep measures | 17.7 ± 0.6 years | Actigraphy | 4 weeks |

| | | | | | | |
|---|---|---|---|---|---|---|
| Sleep quantity and quality is not compromised during planned burn shifts of less than 12 h | 2016 | Deakin University, Geelong, Australia | 33 | | Actigraphy | 4 weeks |
| The impact of long work hours and shift work on cognitive errors in nurses | 2017 | Université de Moncton, Moncton, NB, Canada | 28 | | Actigraphy, sleep questionnaires (KSS) | 4 consecutive shifts (~4 weeks) |
| Sleep and Alertness in Medical Interns and Residents: An Observational Study on the Role of Extended Shifts | 2017 | University of Pennsylvania, Philadelphia, PA, USA | 224 | | Actigraphy | 25 days |
| Examining courses of sleep quality and sleepiness in full 2 weeks on/2 weeks off offshore day shift rotations | 2017 | University of Groningen, Groningen, The Netherlands | 42 | 21–63 years | Actigraphy | 4 weeks |
| Recognizing Academic Performance, Sleep Quality, Stress Level, and Mental Health using Personality Traits, Wearable Sensors and Mobile Phones | 2017 | Massachusetts Institute of Technology, Cambridge, USA | 66 | 20.1 ± 1.5 years | Actigraphy | 30 days |
| Relationships between resting heart rate, heart rate variability and sleep characteristics among female collegiate cross-country athletes | 2018 | University of Connecticut, Korey Stringer Institute, Storrs, CT, USA | 10 | 19 ± 1 years | Actigraphy (WHOOP) | 84 days |
| High heritability of adolescent sleep-wake behavior on free, but not school days: a long-term twin study | 2018 | University of Zurich, Zurich, Switzerland | 51 | 12.8 ± 1.0 years | Actigraphy | 6 months |
| The influence of sleep hygiene education on sleep in professional rugby league athletes | 2018 | Brisbane Broncos Rugby League Club | 24 | 25.4 ± 3.3 years | Actigraphy | 6 weeks |
| Effects of Sleep, Physical Activity, and Shift Work on Daily Mood: A Prospective Mobile Monitoring Study of Medical Interns | 2018 | University of Michigan Medical School, Ann Arbor, MI, USA | 33 | 27.33 ± 2.58 years | Fitbit | 8 months (2 months prior to internship and 6 months after, average 86.5 days per subject) |
| Day-to-day variations in daily rest periods between working days and recovery from fatigue among information technology workers: One-month observational study using a fatigue app | 2018 | National Institute of Occupational Safety and Health, Kawasaki, Japan | 55 | 39.6 ± 6.3 years | Actigraphy | 1 month |
| Sleep, activity and fatigue reported by Postgraduate Year 1 residents: a prospective cohort study comparing the effects of night float versus the traditional overnight on-call system | 2018 | National University Health System, Singapore | 49 (only 11 completed 4 months) | 25-27 years | Actigraphy | 4 months |

| | | | | | | |
|---|---|---|---|---|---|---|
| Dynamics and Ultradian Structure of Human Sleep in Real Life | 2018 | Ludwig Maximilian University Munich, Munich, Germany | 573 | | Actigraphy + 2 night PSG | median 32 days |
| Sleep and Exercise in Emergency Medicine Residents: An Observational Pilot Study Exploring the Utility of Wearable Activity Monitors for Monitoring Wellness | 2018 | University of Saskatchewan, University of Alberta | 20 | | Fitbit | 4 weeks |
| Individual differences in compliance and agreement for sleep logs and wrist actigraphy: A longitudinal study of naturalistic sleep in healthy adults | 2018 | U.S. Army Research Laboratory, UC Santa Barbara | 30 | 18–35 years | Actigraphy, daily sleep diaries | 16 weeks |
| Interindividual and intraindividual variability in adolescent sleep patterns across an entire school term: A pilot study | 2019 | University of South Australia, Adelaide, Australia | 47 | 14-17 years | Actigraphy | 81 days |
| Irregular sleep and event schedules are associated with poorer self-reported well-being in US college students | 2019 | Brigham and Women’s Hospital, Boston, MA, USA | 223 | 19.4 ± 1.5 years | Actigraphy and daily diary | ~ 30 days (6 to 34 days) |
| Effects on resident work hours, sleep duration, and work experience in a randomized order safety trial evaluating resident-physician schedules (ROSTERS) | 2019 | Boston Children’s Hospital; Children’s Hospital Colorado; University of Iowa Stead Family Children’s Hospital; Seattle Children’s Hospital; Cincinnati Children’s Hospital Medical Center; and University of Virginia Children’s Hospital | 302 | | Actigraphy | 1 month |
| Associations of Daily Weather and Ambient Air Pollution with Objectively Assessed Sleep Duration and Fragmentation: A Prospective Cohort Study | 2020 | Beth Israel Deaconess Medical Center, Massachusetts General Hospital, Brigham and Women’s Hospital | 98 | 35.1 ± 12.1 years | Actigraphy and daily questionnaires | At least 6 weeks |
| The Impact of the German Strategy for Containment of Coronavirus SARS-CoV-2 on Training Characteristics, Physical Activity and Sleep of Highly Trained Kayakers and Canoeists: A Retrospective Observational Study | 2020 | University of Würzburg, Würzburg, Germany | 14 | 17.1 ± 1.9 years | Actigraphy | 8 weeks |

| | | | | | | |
|---|---|---|---|---|---|---|
| Examining the Relationship Between Biometric Indicators and Pharmacy Students' Academic Performance | 2020 | Western New England University College of Pharmacy and Health Sciences, Springfield, Massachusetts, USA | 63 | | Fitbit | One semester (January 25, 2016, through May 13, 2016) |
| Continuous 7-Month Internet of Things–Based Monitoring of Health Parameters of Pregnant and Postpartum Women: Prospective Observational Feasibility Study | 2020 | University of Turku, Turku, Finland | 20 | 26 ± 5 years | Garmin watch | 7 months (6 months during pregnancy and 1 month postpartum) |
| Within-person fluctuations in stressful life events, sleep, and anxiety and depression symptoms during adolescence: a multiwave prospective study | 2020 | Harvard University, Cambridge, MA, USA | 30 | 15-17 years | Fitbit | 12 months |
| Multidimensional sleep health is not cross-sectionally or longitudinally associated with adiposity in the Study of Women's Health Across the Nation (SWAN) | 2020 | University of Pittsburgh, Pittsburgh, PA, USA | 221 | 52.1 ± 2.1 years | Actigraphy | Entire menstrual cycle or 35 days |
| Practice parameters for the use of actigraphy in the military operational context: the Walter Reed Army Institute of Research Operational Research Kit Actigraphy (WORK-A) | 2020 | Behavioral Biology Branch, Walter Reed Army Institute of Research, Silver Spring, MD, USA | 286 | | Actigraphy | 31 days |
| Work e-mail after hours and off-job duration and their association with psychological detachment, actigraphic sleep, and saliva cortisol: A 1-month observational study for information technology employees | 2021 | National Institute of Occupational Safety and Health, Kawasaki, Japan | 58 | 39.3 ± 6.2 years | Actigraphy | 1 month |
| Diurnal variations in multi-sensor wearable-derived sleep characteristics in morning- and evening-type shift workers under naturalistic conditions | 2021 | University of Zurich, Zurich, Switzerland | 89 | 33.85 ± 7.73 years | Fitbit | 1 month |
| Contactless Sleep Monitoring for Early Detection of Health Deteriorations in Community-Dwelling Older Adults: Exploratory Study | 2021 | University of Bern, Bern, Switzerland | 37 | 70-101 years | Bed sensor under mattress (EMFIT) | 1 year |
| What time do you plan to sleep tonight? An intense longitudinal study of adolescent daily sleep self-regulation via planning and its associations with sleep opportunity | 2021 | Monash University, Melbourne, Australia | 205 | 16.9 ± 0.9 years | Actigraphy, and daily sleep diaries | 28 days |

| | | | | | | |
|---|---|---|---|---|---|---|
| Trends in Strategic Napping in Surgical Residents by Gender, Postgraduate Year, Work Schedule, and Clinical Rotation | 2021 | MedStar Institutes for Innovation, Washington, DC | 22 | 30.09 ± 2.77 years | Actigraphy (Zulu watch) | 8 weeks |
| Self-reported reasons for on-duty sleepiness among commercial airline pilots | 2021 | Finnish Institute of Occupational Health, Helsinki, Finland | 86 | | Actigraphy | 2 months |
| Tracking Sleep, Temperature, Heart Rate, and Daily Symptoms Across the Menstrual Cycle with the Oura Ring in Healthy Women | 2022 | University of California, Irvine, CA, USA | 26 | 18-35 years | Oura Ring | One complete menstruation cycle |
| Sleep of recruits throughout basic military training and its relationships with stress, recovery, and fatigue | 2022 | Army Recruit Training Centre, Blamey Barracks, Kapooka, Australia | 45 | 25.2 ± 7. 2 years | Actigraphy and daily 5-point Likert scale | 12 weeks |
| Robust stability of melatonin circadian phase, sleep metrics, and chronotype across months in young adults living in real-world settings | 2022 | Brigham and Women's Hospital, Boston, MA, USA | 15 | 19.1 ± 0.3 years | Actigraphy and twice daily sleep diaries | ~ 100 days |
| The submariners' sleep study: a field investigation of sleep and circadian hormones during a 67-day submarine mission with a strict 6-h-on/6-h-off watch routine | 2022 | Royal Military Academy, Brussels, Belgium | 9 | 27.36 ± 3.57 years | Actigraphy and sleep diary | 67 days |
| Sleep and physical activity: results from a long-term actigraphy study in adolescents | 2022 | University of Bern, Bern, Switzerland | 50 | 12.78 ± 1.02 years | Actigraphy | 6 months |
| Use of the Xiaomi Mi Band for sleep monitoring and its influence on the daily life of older people living in a nursing home | 2022 | Universidade da Coruña, Spain | 21 | 86.38 ± 9.26 years | Xiaomi band | 1 year |
| Transition to shift work: Sleep patterns, activity levels, and physiological health of early-career paramedics | 2022 | University of Tasmania, Hobart, Tasmania, Australia | 28 | | Actigraphy | 25 days x 3 (baseline, month 1, month 5) |
| Fluctuations in behavior and affect in college students measured using deep phenotyping | 2022 | Harvard University, Cambridge, MA, USA | 49 | 18-19 years | Actigraphy | One academic year + a few summer days |
| Importance of Getting Enough Sleep and Daily Activity Data to Assess Variability: Longitudinal Observational Study | 2022 | Reykjavík University, Reykjavík, Iceland | 67 | | Actigraphy + mattress sensor | >1 month |
| Assessing the Effect of Extreme Weather on Population Health Using Consumer-Grade Wearables in Rural Burkina Faso: Observational Panel Study | 2023 | Centre de Recherche en Santé, Nouna, Burkina Faso University Hospital, | 143 | | Actigraphy | 11 months |

| | | | | | | |
|---|---|---|---|---|---|---|
| | | Heidelberg University, Heidelberg, Germany | | | | |
| Sleep impairment and altered pattern of circadian biomarkers during a long-term Antarctic summer camp | 2023 | Universidade Federal de Minas Gerais, Belo Horizonte, MG, Brazil | 7 | 32.3 ± 8. 4 years | Actigraphy | 62 days |
| Exploratory study of the effects of sex and hormonal contraceptives on alertness, fatigue, and sleepiness of police officers on rotating shifts | 2023 | McGill University, Montréal, Québec, Canada | 56 men, 20 women | 32.0 ± 5.3 years | Actigraphy and daily questionnaires | Complete work cycle (35 or 28 days) |
| Evaluation of the effectiveness of sleep hygiene education and FITBIT devices on quality of sleep and psychological worry: a pilot quasi-experimental study among first-year college students | 2023 | University of Sharjah, Sharjah, United Arab Emirates | 50 | 17.9 ± 0.29 years | Fitbit | 4 weeks |
| Objective Prediction of Next-Day's Affect Using Multimodal Physiological and Behavioral Data: Algorithm Development and Validation Study | 2023 | University of California, Irvine, Irvine, CA, United States | 20 | 19.80 ± 1.0 year s | Samsung smartwatch and Oura ring | 12 months (range from like 20 days to a year) |
| Country differences in nocturnal sleep variability: Observations from a large-scale, long-term sleep wearable study | 2023 | National University of Singapore, Singapore | 226,187 | | Oura Ring | 52 weeks (minimum 90 days) |
| Good sleep is a mood buffer for young women during menses | 2023 | University of California, Irvine, Irvine, CA, USA | 72 | 18-33 years | Oura Ring, daily sleep diaries | 1 month |
| Nightly sleep duration predicts grade point average in the first year of college | 2023 | Carnegie Mellon University, Pittsburgh, PA, USA | 634 | | Fitbit | 1 month |
| Sleep during COVID-19 Pandemic: A Longitudinal Observational Study Combining Multisensor Data with Questionnaires | 2023 | Computer Science Department Aalto University Helsinki FI | 111 | | Fitness tracker (Polar Ignite) | 1 year |
| Sleep regularity in healthy adolescents: Associations with sleep duration, sleep quality, and mental health | 2023 | University ofBern, Bern, Switzerland | 46 | 12.78 ± 1.07 years | Actigraphy | 6 months |
| Effect of sleep report feedback using information and communication technology combined with health guidance on improving sleep indicators in community-dwelling older people: a pilot trial | 2023 | Osaka University,Osaka, Japan | 29 | | Under mattress activity sleep analyzer | 3 months |

| | | | | | | |
|---|---|---|---|---|---|---|
| Nighttime Ambient Temperature and Sleep in Community-Dwelling Older Adults | 2024 | Marcus Institute for Aging Research, Boston, MA, USA | 50 | > 65 years | Oura Ring | 31–358 days |
| Sleep health and anxiety symptoms in midlife women: the study of women's health across the nation (SWAN) | 2024 | Chicago, Southeast Michigan, Pittsburgh, Oakland (CA) | 270 | 51.8 ± 2. 2 years | Actigraphy | Entire menstrual cycle or 35 days |
| Investigating the Feasibility of Using a Wearable Device to Measure Physiologic Health Data in Emergency Nurses and Residents: Observational Cohort Study | 2024 | University of Pennsylvania, USA | 20 | | Actigraphy (WHOOP) | 6 weeks |
| Trends in sensor-based health metrics during and after pregnancy: descriptive data from the apple women's health study | 2024 | Apple Inc., Cupertino, CA, USA | 733 participants, 757 pregnancies | 29–35 years | Apple watch | 12 weeks prior to last menstrual period to 12 weeks postpartum |
| Interaction of sleep and emotion across the menstrual cycle | 2024 | DeBakey VA Medical Centre, Houston, TX, USA | 51 | 18-35 years | Actigraphy, daily sleep diaries | 2 menstrual cycles |
| The organization of sleep–wake patterns around daily schedules in college students | 2024 | Harvard Medical School, Boston, MA, USA | 223 | 18-27 years | Actigraphy | ~30 days |
| Stability and Volatility of Human Rest-Activity Rhythms: Insights from Very Long Actograms (VLAs) | 2024 | Baylor College of Medicine, Houston, TX USA | 48 | | Actigraphy | More than 90 days |
| The Two Fundamental Shapes of Sleep Heart Rate Dynamics and Their Connection to Mental Health in College Students | 2024 | University of Vermont, Burlington, VT, USA | 603 | | Oura Ring | 8 weeks |
| Sleep, Well-Being, and Cognition in Medical Interns on a Float or Overnight Call Schedule | 2024 | Yong Loo Lin School of Medicine, National University of Singapore, Singapore | 96 | 24.7 ± 1.1 years | Oura Ring | 8 weeks |
| Precision Assessment of Real-World Associations Between Stress and Sleep Duration Using Actigraphy Data Collected Continuously for an Academic Year: Individual-Level Modeling Study | 2024 | Harvard University, Cambridge, MA, USA | 55 | | Actigraphy | Academic year |
| The effects of physical activity on sleep architecture and mood in naturalistic environments | 2024 | The University of Texas at Austin, TX, USA | 65 | 18-35 years | Fitbit | Mean 35.2 days |
